# Increased cocaine self-administration and prefrontal cortical dysregulation of glutamatergic and GABAergic signaling in a rat model for sensory processing sensitivity

**DOI:** 10.1101/2025.10.28.685083

**Authors:** Rick Hesen, Francesca Mottarlini, Lucia Caffino, Sophie Fennema, Susanna Parolaro, Cyprien G. J. Guerrin, Boris B. Quednow, Fabio Fumagalli, Michel M.M. Verheij, Judith R. Homberg

## Abstract

While all organisms are tuned to the environment, some are more sensitive than others. In humans, this is reflected by the trait sensory processing sensitivity (SPS), comprising 20-30% of the population, which is characterized by heightened emotional reactivity, deeper information processing, greater awareness of their environment, and susceptibility to overstimulation. It is expected but unproven that high SPS increases the risk for psychostimulant use. To clarify this, we selected Wistar rats on behavioral extremes related to SPS and subjected them to isolation, neutral, and enriched housing conditions, followed by cocaine self-administration (S/A). Subsequently, we investigated neurobiological differences in excitatory/inhibitory neurotransmission of the low- and high SPS-like rats by evaluating key glutamatergic and GABAergic proteins in the infralimbic (ILc) and prelimbic (PLc) subregions of the prefrontal cortex. High SPS-like rats displayed higher cocaine intake during 1h S/A training sessions in isolated versus social conditions, whereas low SPS-like rats showed similar intake across housing conditions. During 6h S/A sessions, high SPS-like rats displayed a higher cocaine intake regardless of housing and greater motivation for cocaine under a progressive ratio scheme. High SPS-like cocaine-naïve rats exhibited lower expression of glutamatergic and GABAergic markers in ILc and PLc compared to low SPS-like cocaine-naïve rats. The opposite occurred after cocaine exposure: low and high SPS-like rats showed lower and higher expression levels, respectively, compared to naïve counterparts. In conclusion, these findings suggest that a high SPS-like trait with associated glutamatergic and GABAergic alterations might be a risk factor for the development of cocaine use disorder.

## Introduction

In our digitalized postmodern world, we are constantly exposed to a barrage of stimuli ranging from socio-economic pressures and family disruptions to social media overload and distressing global events such as climate change and war. Responses to these stressors vary widely, with individuals high in “sensory processing sensitivity” (SPS) – a personality trait present in 20-30% of the human population [1–5] – being especially susceptible to intense sensory and emotional stimuli. SPS is moderately heritable [6], evolutionarily conserved [7–9], and characterized by four main facets: 1) heightened emotional reactivity, 2) deeper processing of sensory information, 3) greater awareness of subtle environmental stimuli, and 4) enhanced susceptibility to overstimulation [4].

High SPS-individuals tend to flourish under supportive conditions [10–13], but are at risk to perish under aversive ones [13, 14]. Thus, the trait confers both risk and resilience to affective problems, depending on context. Neurobiologically, high SPS individuals display stronger neural activation in response to emotional cues [13, 15], and altered connectivity in regions involved in cognitive- and emotional processing (e.g., limbic system and prefrontal cortex, PFC) [13, 15–17], with the most pronounced alterations in axonal microarchitecture in the (ventromedial) PFC [18]. Together, these findings highlight that SPS is not only a psychological phenotype but also marked by functional and structural differences in the frontal circuitry involved in emotional regulation and cognitive control.

It has been postulated that psychostimulants are used, at least in part, to modulate affective and cognitive states as a functional adaptation to modern environments [19]. Thus, under adverse circumstances, especially when negative urgency is in play [20], some individuals start to use psychostimulants as coping strategy [21], which can escalate into compulsive use or a substance use disorder (SUD). Consistently, environmental adversity is a known risk factor for SUD development [22–24] and enhances psychostimulant preference in preclinical models [25–27]. Intrinsic traits further modulate risk: higher trait anxiety, closely linked to emotional reactivity of SPS [28, 29], is associated with greater psychostimulant use [30–33]. In addition, developmental sensory overstimulation increases the vulnerability to cocaine later in life [34]. Thus, both environmental adversity and SPS-related traits may converge to increase vulnerability to psychostimulant use.

Neurally, psychostimulants modulate the glutamate homeostasis in the PFC [35], a region that is implicated in high SPS. More specifically, chronic psychostimulant use alters glutamine/glutamate in frontal white matter, anterior cingulate cortex and the PFC [36, 37]. These changes are corroborated by animal models where low dosages of methamphetamine or cocaine shift glutamine/glutamate and glutamate/GABA ratios in the PFC [38, 39]. While no such data are available for SPS, high SPS has been associated with increased absolute power of PFC neural oscillations, pointing to higher activity of cortical excitatory cells [40]. We postulate that high SPS individuals exhibit increased neural activity, which renders them more sensitive to environmental stimulation but simultaneously reduces the dynamic range for adaptation to overstimulation due to a ceiling effect [41]. As a functional adaptation, psychostimulant use may serve to increase this dynamic range.

In this study, we aimed to test the hypothesis that high SPS, particularly under adverse environments, increases risk of psychostimulant use and that this relates to altered glutamate and GABA signaling in the PFC. As it is hard to capture humans before psychostimulant use to assess SPS level and brain glutamatergic and GABAergic signaling, we used animals for hypothesis testing. First, we selected behavioral extremes associated with SPS in a population of outbred wild-type Wistar rats. Based on the above mentioned core SPS characteristics, aligning with the features of the highly sensitive person scale [42], we used three emotionality-based behavioral assays: 1) the elevated plus maze (EPM) to measure emotional reactivity, 2) conditioned freezing (CF) to assess depth of sensory information processing, and 3) prepulse inhibition (PPI) to measure awareness for environmental subtleties [41]. Rats in the top and bottom 20^th^ percentile, aligned with SPS prevalence estimates in humans [1–4], were classified as high- or low SPS-like, respectively. These rats were housed under isolated (negative), neutral or enriched (positive) conditions to explore environmental sensitivity. Next, they underwent intravenous cocaine S/A training (1h/day), followed by long access (LgA, 6h/day) sessions. Finally, we quantified glutamatergic- and GABAergic proteins in the prelimbic- (PLc) and infralimbic (ILc) cortical subregions of the medial PFC – areas implicated in both SPS [13, 15, 18] and cocaine-seeking behavior [43–47].

## Methods

A detailed description of the experimental methods and statistical analysis can be found in the supplementary methods. We have behaviorally assessed 165 outbred male Wistar rats (56-70 days old) using the EPM (total time (s) in open arms in 5 minutes [48]), CF (mean total freezing time (s) in 1^st^ and every 4^th^ trial out of 24 trials [49]), and PPI (120 dB startle amplitude (P120S) and PPI [50, 51]) tests in two experimental batches (N_1_=55 and N_2_=110). Within each batch, rats were ranked based on their composite scores across all tests (equal weighting per test). The top- and bottom 20% (+2% reserve) were assigned to high- and low SPS-like groups, respectively (Fig.1), and distributed over different housing conditions: isolated- (type II cage, no shelter), neutral- (pair-housed, shelter), or social-housing (6 rats with multiple shelter, bedding material and toys). All high- and low-SPS-like rats received right jugular vein cannulation surgery. Cannulas were flushed daily with heparinized saline (50 USP, LEO) during the entire experiment to prevent blockade. The control group, later referred to as cocaine-naïve rats, also received jugular vein cannulation and were placed in the operant chamber but were not able to obtain cocaine. The cocaine-group was trained to self-administer cocaine; 0.5 mg/kg/infusion, at a fixed ratio 1 (FR1) schedule during daily 1-hour sessions for at least 11 days [52, 53]. Following training, cocaine S/A sessions were extended to long-access (LgA); 6-hour per day on FR1 [52–54]. After 21 sessions and successful escalation under LgA conditions, motivation to work for cocaine was tested under a progressive ratio (PR) schedule of reinforcement [52, 53]. After PR, rats were abstinent from cocaine for two weeks before LgA cocaine S/A was re-established for 7 days (Fig.2). Afterwards, the rats were sacrificed by decapitation and brains were flash frozen on dry ice. The PLc and ILc subregions were punched (Bregma +4.20mm to +2.52mm, Rat Brain Atlas [55]), and left- and right hemisphere samples were pooled and homogenized. Proteins were quantified and analyzed using Western blots for glutamatergic- and GABAergic proteins and normalized using β-actin (43 kDa) loading control (Fig.3). Gels were run in duplicate and reported values represent the average of two independent runs. We used a correction factor for the second gel. Correction factor gel 2 = average of (OD protein of interest/OD β-actin for each sample loaded in gel 1)/(OD protein of interest/OD β-actin for the same sample loaded in gel 2) [56]. Behavioral and molecular data were analyzed using IBM SPSS, RStudio, and GraphPad Prism. Behavioral measures (EPM, CF, P120s, and PPI) were first tested for normality and homoscedasticity, and due to assumption violations, analyzed using Brown-Forsythe-corrected ANOVA with Dunnett T3 post hoc tests. Dimension reduction was performed via robust linear discriminant analysis (rLDA) to account for unequal variances and non-normal data distributions. Cocaine self-administration data were analyzed with linear mixed models to accommodate repeated measures and random effects, while progressive ratio data were analyzed using two-way ANOVA. Molecular data were analyzed with two-way ANOVA (SPS-like status × cocaine status), followed by Benjamini-Hochberg FDR correction and Tukey’s post hoc tests.

**Figure 1.**
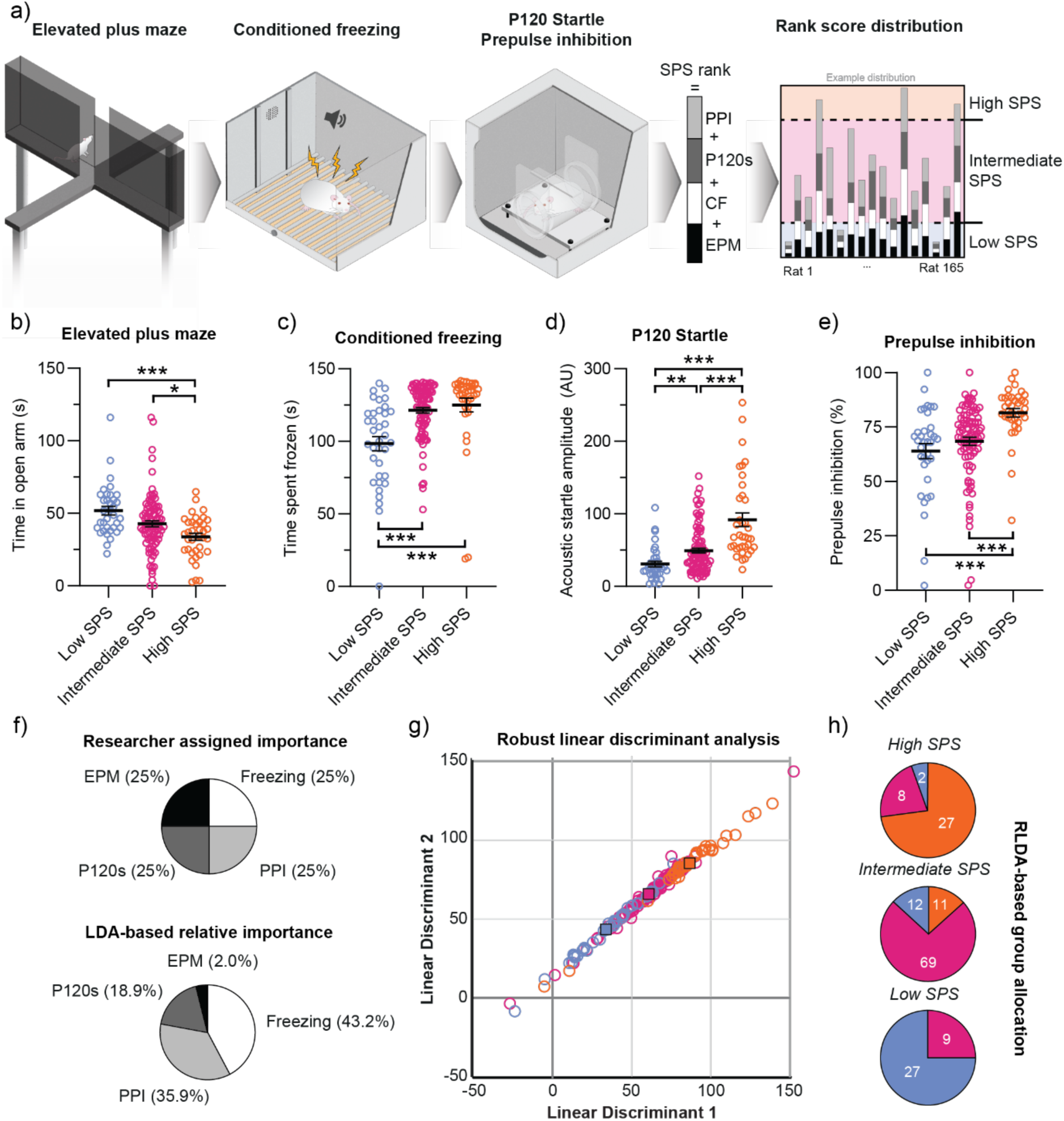
a) Graphical overview of the three behavioral test setups and SPS-rank allocation based on the four outcome variables (EPM, CF, P120s and PPI scores). b-e) Behavioral outcome per SPS with time in open arms in the EPM test (b), freezing time in the CF test (c), startle amplitude after 120 dB pulse in the PPI test (d), and % prepulse inhibition in the PPI test (e). The relative-importance per outcome variable following the researcher-assigned relative importance in comparison to the rLDA-based relative importance (f). Sample distribution along the first two discriminants (LD1, LD2) in a structure matrix derived from the rLDA (g). Reclassification of originally assigned groups (top: High SPS, middle: Intermediate SPS, bottom: Low SPS). Each pie chart shows the proportion of rats that would fall into a different group if classified using the rLDA-derived relative importance rather than the researcher-assigned weighting (h). Number of rats per group are: N_low SPS_=36, N_intermediate SPS_=92, N_high SPS_=37. Data represent group means ± S.E.M. Statistics display (Brown-Forsythe-corrected ANOVA with) Dunnett’s T3-corrected post hoc results: \**p*<0.05, \*\**p*<0.01, \*\*\**p*<0.001.

**Figure 2.**
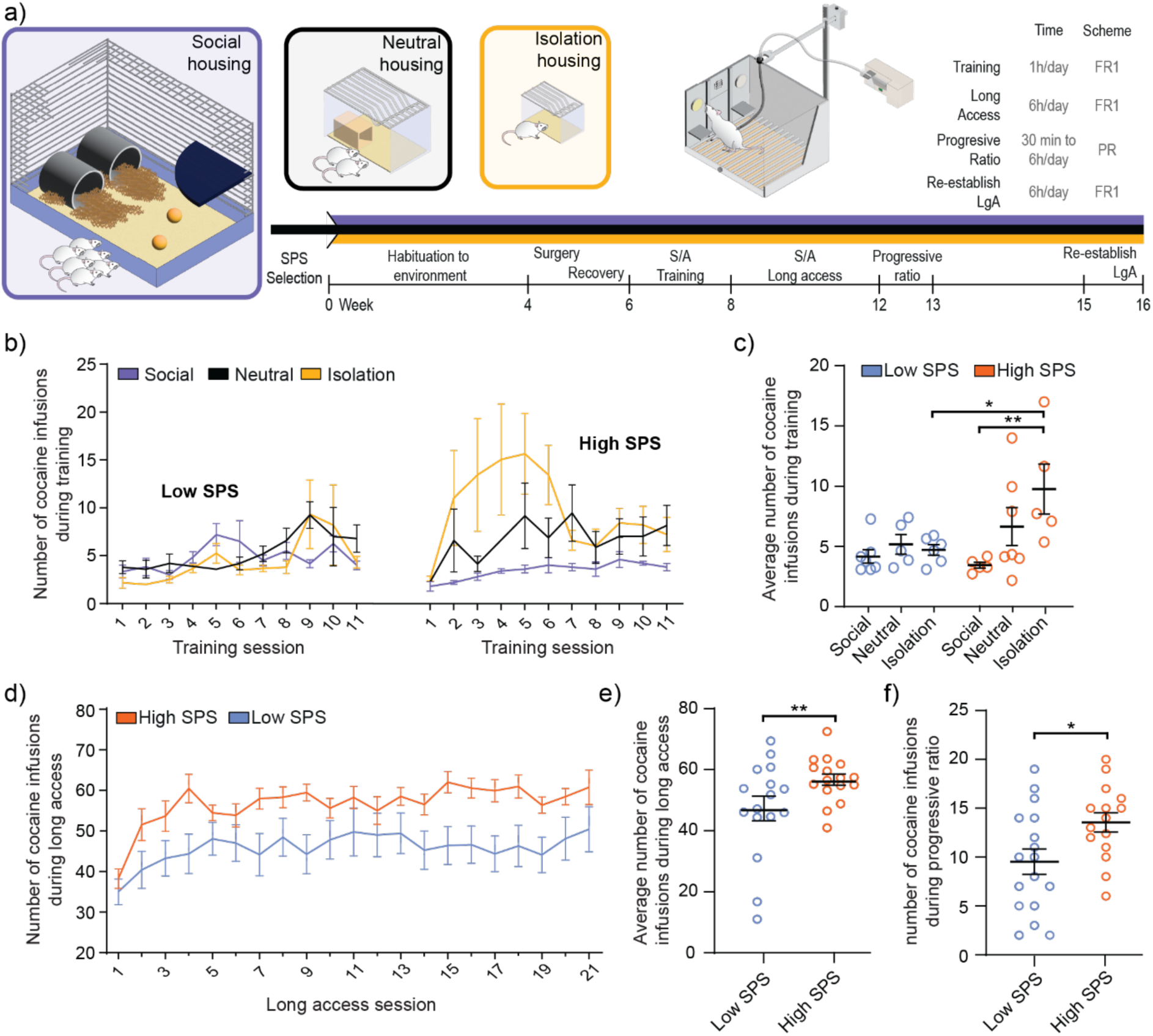
a) Graphical overview of the experimental timeline for housing condition exposure and cocaine self-administration parameters. b) Number of cocaine-infusions during 11 days of S/A training excluding manual shaping. c) Average number of cocaine infusions during the training session per housing condition per SPS. d) Number of self-administered cocaine infusions during 21 days of LgA. e) Average number of cocaine infusions during LgA per SPS group. f) Average number of cocaine infusions during PR. The number of rats per group are: N_low SPS - social_ = 6, N_low SPS - neutral_ = 5, N_low SPS - isolation_ = 6, N_high SPS - social_ = 5_Training_/4_LgA_, N_high SPS - neutral_ = 7, N_high SPS - isolation_ = Data represent group means ± S.E.M. Statistics display Bonferroni-corrected linear mixed model results for 2b-e and two-way ANOVA for 2f with \**p*<0.05, \*\**p*<0.01.

**Figure 3.**
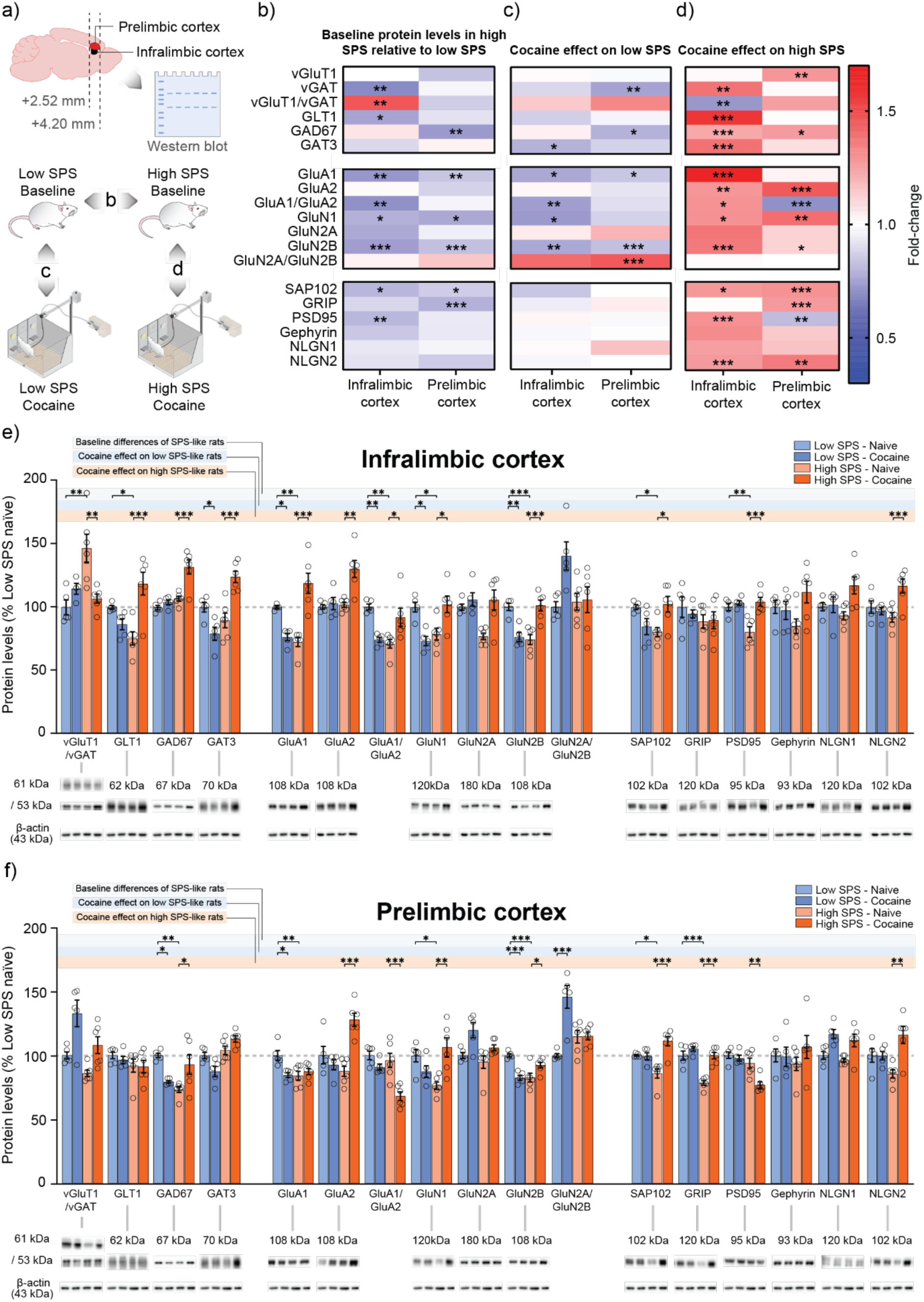
a) Graphical representation of infralimbic- and prelimbic cortical regions and punching coordinates used in the western blot analyses (above) along with group comparison indication for b-d (below). Relative protein expression levels between b) cocaine-naive high SPS-like rats relative to cocaine-naive low SPS-like rats, c) cocaine-exposed low SPS-like rats relative to cocaine-naive low SPS-like rats, and d) cocaine-exposed high SPS-like rats relative to cocaine-naive low high SPS-like rats. Protein expression levels of glutamatergic and GABAergic markers measured in the whole homogenate of the infralimbic (e) and prelimbic (f) cortices are expressed in bar scatter plots as percentage of the mean ± SEM compared to the low SPS-like naive group with representative immunoblots below each subset. The number of rats per group are: N_low SPS - naive_ = 5, N_low SPS – Cocaine S/A_ = 5, N_high SPS - naive_ = 6, N_high SPS – Cocaine S/A_ = 6. Data represent group means ± S.E.M. Statistics display Benjamini-Hochberg corrected two-way ANOVA results with Tukey’s-corrected post hoc tests for 3b-f with \**p*<0.05, \*\**p*<0.01, \*\*\**p*<0.001.

## Results

### Outbred Wistar rats can be classified as high- and low SPS-like based on behavioral readout

We have selected outbred Wistar rats across two batches (N_1_=55 and N_2_=110) based on their compound behavior scores in the EPM, CF, and PPI tests (Fig.1a). We found significant group-level differences for all four behavioral variables (Brown-Forsyth-corrected ANOVA; EPM: *F*_2, 159_ = 7.20, *p*=≤0.001; CF: *F*_2, 87_ = 11.52, *p*≤0.001; P120s: *F*_2, 64_ = 22.04, *p*≤0.001; PPI: *F*_2, 96_ = 7.87, *p*≤0.001, Fig.1b-e). Post-hoc comparisons show that high SPS-like rats scored different in all individual tests compared to their low- (High vs. Low; EPM: *p*≤0.001, CF: *p*≤0.001, P120s: *p*≤0.001, PPI: *p*≤0.001) and intermediate SPS-like counterparts (High vs. Intermediate; EPM: *p*=0.016, P120s: *p*≤0.001, PPI: *p*≤0.001), except in the CF test (High vs. Intermediate; CF: *p*=0.843). The low SPS-like rats scored differently from intermediate SPS-like rats in the CF and P120s tests (Low vs. Intermediate; CF: *p*≤0.001, P120s: *p*=0.004), but not in the EPM and PPI tests (Low vs. Intermediate; EPM: *p*=0.405, PPI: *p*=0.431; Fig.1b-e; for all statistics, Table S1-4).

While all four behavioral measures were weighted equally in the composite SPS-like score, we assessed post-hoc their relative discriminative power using a robust linear discriminant analysis (rLDA) [57, 58]. This analysis revealed that CF (43.2%) and PPI (35.9%) contributed most to group separation, whereas P120 (18.9%) and EPM (2.0%) played smaller roles (Fig.1f). The first two linear discriminants (LD1: 58.1%, LD2: 23.7%) explained 81.8% of the total variance and contributed to a spectrum-based, rather than group separation based, distribution (Fig.1g). The rLDA model achieved an overall classification accuracy of 74.5%, with group accuracies of 75.0% in the low- and intermediate SPS-like groups and 73.0% in the high SPS-like group compared to the researcher assigned group compositions (Fig.1h; for all statistics, Table S5-7).

### High SPS-like rats display increased cocaine intake and motivation, which is affected by housing during S/A training

Based on researcher-assigned importance SPS-like classification, high- and low SPS-like rats were introduced to their novel housing conditions and subjected to cocaine S/A (Fig.2a). These results have been visually compared to rLDA-based classification (Fig.S4). Using linear mixed models, we found that cocaine intake during S/A training was significantly affected by the interactions between SPS and housing (*F*_2, 47.89_ = 4.19, *p*=0.021, Fig.2c) and SPS and time (*F*_10, 102.86_ = 2.38, *p*=0.014; for all statistics, Table S8-13). Post hoc comparison indicated that high SPS-like rats housed in isolated conditions had a significantly higher cocaine intake compared to high SPS-like rats in social-enriched conditions (t_47_ = 4.20, *p*=0.001; Fig.2c), as well as compared to low SPS-like rats in isolation (t_47_ = 3.44, *p*=0.016; Fig.2c; for details, Fig.S1).

During LgA sessions, cocaine intake increased in all rats (time effect: *F*_20, 192.00_ = 4.81, *p*<0.001). Overall, high SPS-like rats exhibited higher cocaine intake (*F*_1, 40.81_ = 9.40, *p*=0.004) independent of housing conditions (Fig.2d, e; for details, Fig.S2-3; for all statistics, Table S14-17). The motivation to obtain cocaine, assessed using a PR schedule, was also significantly higher in high SPS-like rats, as indicated by their elevated breakpoint compared to low SPS-like counterparts (*F*_1, 26_ = 6.21, *p*=0.019; Fig.2f; for all statistics, Table S18-20).

After two weeks of abstinence, LgA cocaine S/A was re-established for 7 days prior to sacrifice, during which neither SPS-like group nor housing conditions had a significant main effect on cocaine intake (Fig.S5).

### High- and low SPS-like rats display differential glutamate- and GABAergic protein expression levels at baseline and after cocaine exposure in the ILc and PLc

Given the absence of housing effects on cocaine intake during LgA and PR sessions, we focused on the impact of cocaine exposure on SPS-like molecular phenotype under neutral housing conditions. We compared glutamatergic and GABAergic signaling-related proteins in the ILc and PLc in cocaine-naïve and cocaine-exposed low and high SPS-like rats (Fig.3a). In this explorative approach, we grouped proteins into three functional categories: (1) presynaptic markers (vGluT1, vGAT, GLT1, GAT3, GAD67), (2) postsynaptic receptor subunits (GluA1, GluA2, GluN1, GluN2A, GluN2B), and (3) scaffolding/adhesion molecules (SAP102, GRIP, PSD95, Gephyrin, NLGN1, NLGN2).

To visualize group-wise expression profiles for both cortices, we generated fold-change heatmaps summarizing relative differences between paired groups (Fig.3a-d). Under cocaine-naïve (baseline) conditions, high SPS-like rats displayed a consistent downregulation of multiple single proteins relative to low SPS-like controls, without significant increases (Fig.3b). Specifically, seven proteins were downregulated in the ILc (vGAT, p=0.001; GLT1, p=0.049; GluA1, p=0.006; GluN1, p=0.037; GluN2B, p<0.001; SAP102, p=0.049; PSD95, p=0.003; Fig.3e, grey row) and six in the PLc (GAD67, p=0.003; GluA1, p=0.008; GluN1, p=0.028; GluN2B, p<0.001; SAP102, p=0.021; GRIP, p<0.001; Fig.3f, grey row). These changes led to an increased vGluT1/vGAT ratio (p=0.002; Fig.3e) and a decreased GluA1/GluA2 ratio (p=0.001; Fig.3e) in the ILc.

Interestingly, cocaine self-administration produced divergent effects depending on SPS-like phenotype. In low SPS-like rats, cocaine exposure led to a downregulation of numerous single proteins in both the ILc and PLc, again without significant increases (Fig.3c). More specifically, we found a reduction in four ILc proteins (GAT3, p=0.0497; GluA1, p=0.025; GluN1, p=0.010; GluN2B, p=0.003; Fig.3e, blue row) and four PLc proteins (vGAT, p=0.003; GAD67, p=0.029; GluA1, p=0.014; GluN2B, p<0.001; Fig.3f, blue row). These changes resulted in a decreased GluA1/GluA2 ratio in the ILc (p=0.006; Fig.3e) and an increased GluN2A/GluN2B ratio in the PLc (p<0.001; Fig.3f).

Conversely, in high SPS-like rats, cocaine exposure reversed the previously observed downregulations (Fig.3d). Eleven single proteins were significantly upregulated in the ILC (vGAT, p=0.003; GLT1, p<0.001; GAD67, p<0.001; GAT3, p<0.001; GluA1, p<0.001; GluA2, p=0.001; GluN1, p=0.014; GluN2B, p<0.001; SAP102, p=0.019; PSD95, p<0.001; NLGN2, p=0.003; Fig.3e, orange row) and eight in the PLc (vGluT1, p=0.01; GAD67, p=0.022; GluA2, p<0.001; GluN1, p=0.003; GluN2B, p=0.044; SAP102, p<0.001; GRIP, p<0.001; NLGN2, p=0.001; Fig.3f, orange row). Notably, no proteins were significantly decreased in the ILc, and only 1 (PSD95) was reduced in the PLc (p=0.002; Fig.3f). These changes resulted in a decreased vGluT1/vGAT ratio (p=0.004, Fig.3e) and increased GluA1/GluA2 ratio (p=0.014; Fig.3e) in the ILc, but a reduced GluA1/GluA2 ratio in the PLc (p<0.001; Fig.3f).

In summary, we found a broad downregulation in GABA- and glutamatergic markers in high SPS-like rats under naïve conditions (Fig.3b), whereas post-cocaine changes were different between groups: low SPS-like rats exhibited a general downregulation GABA- and glutamatergic markers (Fig.3c), whereas a widespread upregulation was seen in high SPS-like rats (Fig.3d).

## Discussion

In this study, we have established an SPS-like rat model through a behavioral selection procedure designed to capture individual variability in a healthy population. Our data revealed that high SPS-like rats displayed increased vulnerability to cocaine seeking behavior. More specifically, high SPS-like rats exhibited greater housing-dependent cocaine intake during short access (training), and housing-independent intake during long access, as well as a higher motivation to obtain cocaine during PR. The housing effect observed during short access sessions underscores the environmental sensitivity of this model. This behavioral profile was paralleled by distinct glutamatergic and GABAergic protein expression profiles in the ILc and PLc. Specifically, high SPS-like rats exhibited a potential presynaptic excitatory state due to reduced inhibition combined with a reduced postsynaptic glutamate receptor availability at baseline. After a history of cocaine S/A, the molecular profile turned to an upregulation of both glutamatergic and GABAergic synaptic proteins, opposite to the downregulation of these proteins in low SPS-like rats.

Using the three behavioral assays aligning with the merely negative valanced features of the highly sensitive person scale [42], we developed a composite score to rank SPS-like traits. Using a post-hoc rLDA we cross-validated the classification accuracy and found an accuracy of 75% across two independent cohorts, exceeding chance levels. The dimension reduction across the first two discriminants resulted in an organization of rats along a near-linear spectrum, aligning with the idea that SPS represents a continuous trait rather than low, intermediate and high SPS define discrete categories [4, 41, 59]. However, the classification was mainly driven by the PPI and CF tests, whereas the contribution of the EPM test was nearly neglectable. A plausible explanation is that the PPI and CF tests measure primarily immediate, stimulus-driven responses that occur automatically in reaction to specific sensory cues, whereas EPM behavior involves active exploration and voluntary decision-making under conflict or uncertainty. In support, PPI is regarded as a mere reflexive sensorimotor gating response and in both rodents and humans modulated by cortical top-down control from the forebrain [60–62]. Notably, motivationally salient prepulses can either reduce or enhance PPI, based on whether they are predicting reward [63] or threat [64], respectively. Related to the latter, the threat expectation induced by looming results in an active, adaptive sensorimotor gating mechanism that helps the organism engage with relevant environmental cues, enhancing chances of survival in complex settings [62]. Similarly, conditioned freezing during extinction in the CF test is well-known to be modulated by prefrontal control over the amygdala in both rodents and humans [65]. CF, and particularly its duration, reflects active sensory integration rather than a simple fear reflex [66, 67]. During sustained freezing, sensory intake is enhanced [68] and information processing is optimized via top-down control from the dorsal anterior cingulate cortex promoting adaptive threat processing [66]. In contrast, for the EPM test there are no clearly defined cross-species neural mechanisms. EPM-related behavior might reflect a more acute, fear-based response in an anxiogenic context rather the more nuanced, trait-like emotionality sensitivity as seen in SPS, such as prolonged affective processing [2, 13]. This could explain its weaker predictive contribution. The limited predictive power of the EPM test suggests that SPS-related emotionality may be better captured by other tasks. Future studies could therefore compare EPM with alternative paradigms, including simple social-choice (or affective bias tasks), which would allow assessment of the positive-affective dimensions of SPS, such as enhanced engagement with rewarding stimuli.

Still, using our behavioral assessment, we observed transient environmental sensitivity in relation to cocaine S/A during training in high SPS-like rats. As expected, high SPS-like rats displayed significantly higher cocaine intake under aversive (isolated) conditions compared to both socially housed counterparts as well as low SPS-like rats under identical, isolated conditions. The finding that LgA cocaine intake and PR responding was higher in high SPS-like rats regardless of housing conditions suggests a shift from a potential environmental modulation to a trait-driven vulnerability, specifically an escalation of cocaine intake and increased motivation to self-administer the drug, respectively. Importantly, the pattern of effects was preserved when SPS grouping was based on the rLDA classification (Fig.s4), although statistical significance was not reached in this reduced dataset. This pattern suggests that environmental enrichment might buffer effects that are lost under prolonged drug exposure. These findings are consistent with the idea that environmental enrichment can buffer cocaine intake, although these effects are dose- and sex-dependent. Protective effects have been reported at 0.1 mg/kg/infusion but not at the higher 0.5 mg/kg/infusion used here [69], and between paired (neutral) and isolated conditions in females but not males [70], which may partly explain the limited impact of housing in our male-only study. Additionally, loss of enrichment has been shown to enhance cocaine reward relative to consistent standard housing [26], potentially contributing to the absence of housing effects during LgA, when all rats were isolated for six hours during their active phase. After abstinence, cocaine intake during the seven days of re-establishment of LgA converged towards low SPS-like levels along with greater variability within groups (Fig.S5d). The loss of SPS effect on cocaine intake might be explained by withdrawal effects experienced during abstinence, but could also reflect a progressive, time-dependent functional adaptation in high SPS-like rats over the course of cocaine exposure. Early in training, high SPS-like rats display both heightened intake and environmental sensitivity, but prolonged exposure appears to trigger neuroplastic changed, supported by the opposing molecular findings in the ILc and PLc, which in turn may normalize the behavior by reducing first environmental sensitivity and afterwards overall heightened vulnerability. However, this proposed dynamic functional adaptation remains speculative and should be tested in a follow-up study that capture neural dynamics across the course of cocaine exposure. In summary, high SPS-like rats display increase cocaine intake and environmental sensitivity during early drug use. During prolonged cocaine use, the environmental sensitivity is lost and after abstinence, cocaine intake behavior normalizes.

Delving deeper into these neurobiological changes, our molecular analysis revealed that high SPS-like rats displayed a distinct glutamatergic/GABAergic protein profile in the ILc and PLc. At baseline, these rats showed a reduction of inhibitory mechanisms in the presynaptic compartment, alongside reduced post-synaptic AMPA- and NMDA receptor availability. Following cocaine-exposure, opposing changes were found in the high- and low SPS-like rats, with more pronounced changes in high SPS-like rats. Together, the findings suggest that the increased cocaine self-administration and motivation in high SPS-like rats is accompanied by excitatory-inhibitory dysregulation in the mPFC.

Specifically, in the ILc of naïve high SPS-like rats, the combination of reduced vGAT and GLT1, and increased vGlut1/vGAT ratio suggests an elevated glutamatergic tone by reduced inhibition. However, this suggested increased excitatory neurotransmission seems accompanied by a compensatory reduction in the expression of the postsynaptic glutamate receptor subunits GluA1, GluN1 and GluN2B, along with SAP102, a GluN2B-related scaffolding protein, and PSD95, an index of synaptic integrity. These findings point to a hypo-reactive glutamatergic response at post-synaptic level. A similar outcome, though via different mechanisms, was observed in the PLc. Here, reduced GAD67 expression suggests a lower GABA production and potential reduced GABAergic tone. Post-synaptically, the overall downregulation of GluA1, GluN1, GluN2B glutamate receptor subunits and their associated scaffolding proteins, SAP102 and GRIP, are again indicative of a hypo-reactive glutamatergic response. These changes align with the concept that high SPS-individuals exhibit increased levels of neuronal activity [40] due to reduced GABAergic inhibition, along with a less responsive glutamatergic post-synapse, potentially indicative for a lower capacity to dynamic adaptation [41].

Following cocaine S/A, these systems diverged further. In low SPS-like rats, changes were relatively modest, while a more profound neuroplastic shift occurred in high SPS-like rats. In the ILc of the latter, the predicted presynaptic upregulation of the GABA production, release, and reuptake (i.e., GAD67, vGAT, GAT3), and the glutamatergic clearance (GLT1) along with the increased post-synaptic receptor subunits expression (i.e., GluA1, GluA2, GluN1, GluN2B) and associated scaffolding proteins (i.e., SAP102, PSD95 and NLGN2) may imply a compensatory response to regulate excitability after cocaine. This may increase the dynamic range for dealing with environmental challenges under high SPS-like conditions. Notably, the increase of GluA1, and its consequential significant increase of the GluA1/A2-ratio, indicates insertion GluA2-lacking Ca^2+^-permeable AMPA (CP-AMPA) receptors [71], which is associated with increased excitability and excitotoxic vulnerability [72]. Alongside increased GluN2B, this may shift the molecular composition of the synapse towards a developmentally plastic state [73], potentially fostering drug craving incubation [74]. These changes were largely opposite in low SPS-like rats, consistent with an overall reduced glutamatergic transmission.

In the PLc, high SPS-like rats with a history of cocaine S/A exhibited increased excitatory and inhibitory presynaptic (i.e. vGluT1 and GAD67) proteins as well as increased post synaptic glutamatergic (i.e., GluA2, GluN1 and GluN2B) determinants. Notably, the selective reduction of the GluA1/A2 ratio, opposite to the ILc, suggests the insertion of GluA2-containing Ca^2+^-impermeable AMPA receptors, potentially indicative for frontal hypoactivity. This hypoactivity has been linked to punishment-resistant cocaine-seeking [75]. Relatedly, targeted magnetic stimulation of PFC or optogenetic stimulation of PLc reduces craving and cocaine seeking in clinical [76, 77] as well as preclinical [75, 78] studies. In contrast, low SPS-like rats showed attenuated responses to cocaine. Cocaine reduced presynaptic GABA synthesis and release (GAD67 and vGAT) and post-synaptic glutamatergic receptors (GluA1 and GluN2B) in the PLc, highlighting an opposite mechanism to that observed in high SPS-like rats. These findings corroborate the large body of evidence of cortical excitation-inhibition imbalances commonly associated with substance use in clinical- and preclinical research [79–84]. The evident opposing effects of cocaine on prefrontal protein expression – an increase in high SPS-like rats and decrease in low SPS-like rats – warrant further investigation.

Our data come with some limitations. First, because men are more vulnerable to cocaine use disorder than women [85], we have tested male rats only. However, our findings may not generalize to female rats. Second, the current behavioral test battery to select SPS-like animals focuses solely on responses to aversive or negative stimuli. The cocaine intake and associated GABAergic and glutamatergic plasticity in SPS-like animals selected based on their behavioral response to rewarding or positive stimuli remains to be further investigated. Third, our molecular approach is explorative and does not establish causality.

In conclusion, we have proposed a behavioral selection approach based on key facets of SPS to model high- and low SPS-like phenotypes in rats. The high SPS-like rats displayed increased environmental sensitivity during acquisition of cocaine S/A, and overall higher cocaine intake during LgA as well as higher breakpoint in PR responding independent of housing conditions. Neurobiologically, this was associated with a baseline excitatory-inhibitory imbalance in the ILc and PLc, indicative of reduced presynaptic inhibition and reduced postsynaptic glutamate receptor expression. Cocaine S/A induced opposing neuroplastic responses, namely a pronounced upregulation of excitatory and inhibitory markers in high SPS-like rats versus a dampening in low SPS-like rats. This pioneering work sets the stage for further understanding of the neurobiological and molecular underpinnings of environmental sensitivity and individual differences in susceptibility to cocaine addiction.

## Data availability statement

Data are available upon request from the corresponding author. The individual datapoints and full statistics are presented in the supplementary file.

## Acknowledgements

We thank the animal caretakers of the Radboud University Medical Center animal facility for their excellent support with animal husbandry.

## Author contributions

JH, FF and BBQ acquired funding. JH and MV conceived the study design. MV, RH, SF and FM acquired the data. RH and FM analyzed the data. RH wrote the draft manuscript and created the figures. FM, LC, BBQ, FF, MV and JH contributed to data interpretation and critically reviewed and edited the manuscript. All authors provided final approval for the manuscript.

