## supplementary methods for "Increased cocaine self-administration and prefrontal cortical dysregulation of glutamatergic and GABAergic signaling in a rat model for sensory processing sensitivity"

Animal details

SPS rank selection – Elevated plus maze, conditioned freezing and prepulse inhibition

Housing conditions

Intravenous catheterization

Cocaine self-administration

Tissue collection and protein isolation

Western blot analysis

Statistical analysis

### Supplementary Code

Robust Linear Discriminant Analysis code for RStudio

### Supplementary Tables

Table S1. SPS selection criteria Levene's test homogeneity of variances test results

Table S2. SPS selection criteria Shapiro-Wilk normality test results

Table S3. SPS selection criteria Brown-Forsythe corrected one-way ANOVA test results

Table S4. SPS selection criteria Dunnett T3 post-hoc results

Table S5. Group-specific constants and group means per SPS-selection criteria using in the rLDA

Table S6. Within-groups covariance matrix used in the rLDA

Table S7. Linear discriminant coefficients used in the rLDA.

Table S8. LMM model dimensions for cocaine S/A training

Table S9. LMM information criteria for cocaine S/A training

Table S10. LMM Type III test of fixed effects for cocaine S/A training

Table S11. LMM estimated marginal means and pairwise comparison of SPS effect in cocaine S/A training

Table S12. LMM estimated marginal means and pairwise comparison of housing effect in cocaine S/A training

Table S13. LMM estimated marginal means and pairwise comparison of SPS*housing interaction effect in cocaine S/A training

Table S14. LMM model dimensions for cocaine S/A long access

Table S15. LMM information criteria for cocaine S/A long access

Table S16. LMM Type III test of fixed effects for cocaine S/A long access

Table S17. LMM estimated marginal means and pairwise comparison of SPS effect in cocaine S/A long access

Table S18. Progressive ratio Levene’s test homogeneity of variances test results

Table S19. Progressive ratio Shapiro-Wilk normality test results

Table S20. Progressive ratio two-way ANOVA test results for SPS, housing and SPS*housing

Table S21. Western blot antibody table

Table S22. Western blot ANOVA results

Table S23. Benjamini-Hochberg corrected Western blot ANOVA results

**Supplementary Figures**

Figure S1. Line chart with number of cocaine infusions per rat during S/A training per SPS and housing

Figure S2. Line chart with number of cocaine infusions per rat during S/A LgA per SPS and housing

Figure S3. Long access cocaine S/A results for housing*SPS and housing

Figure S4. Average number of cocaine infusions during training and LgA with rLDA-based groups

Figure S5. Average number of cocaine infusions during reinstatement for housing, SPS and housing*SPS

Figure S6. Western blot cropped immunoblots for infralimbic cortex

Figure S7. Western blot cropped immunoblots for prelimbic cortex

Figure S8. Western blot protein levels per individual protein for infralimbic cortex

Figure S9. Western blot protein levels per individual protein for prelimbic cortex

**Supplementary References**

### Supplementary Methods

#### *Animals*

A total of 165 outbred adult male Wistar rats (56-70 days old; Crl:WI(WU) // Charles River) were obtained in two experimental batches (N=55 and N=110) from Charles River Laboratories (Germany). The rats were pair-housed (Conventional type III rat cage) and acclimatized for two weeks under reversed light-dark cycle (lights on phase: 20:00-08:00, light off phase (dim red light: 08:00-20:00) in a temperature- (21 ± 1 °C) and humidity- (55 ± 5%) controlled room. They had ad libitum access to water and food, except during testing. Prior to all behavioral tests, rats were habituated in a pre-testing room for one hour. All experiments were approved by the Central Committee Animal Experiments in The Hague, The Netherlands (AVD1030020174387). All efforts were made to minimalize animal suffering and to reduce the number of rats used.

***SPS rank selection***

An SPS-like scale was used based on four outcome variables; time in open arms in the elevated plus maze (EPM) test, freezing time in conditioned freezing (CF) test, and maximum startle amplitude at 120 dB pulse (P120) and the average prepulse inhibition (PPI, average of $PPI_{avg}$ and $PPI_{Vmax}$) in the PPI test. For each test, the rats received a rank score relative to the performance of the group. After all tests, the SPS rank was based on the average rank score of all test ranks with equal weight per test variable. The 20% highest and lowest rank scores were labelled as high- and low SPS-like, and 2% reserves were taken along in case of drop-out during the follow-up procedures.

The EPM [1] consisted of four perpendicularly oriented arms: two open (50 cm x 10 cm) and two enclosed (50 cm x 10 cm x 40 cm) arms, constructed from black polymethyl methacrylate (PMMA). The maze was elevated 50 cm above the floor with room illumination maintained at 10 lux. Rats were allowed to freely explore the maze for five minutes while their movements were recorded [1]. The time (seconds) in each testing arm was quantified automatically using Ethovision XT version 10 [2]. The cumulative time spent in both open arms was used to determine SPS-rank for EPM.

The CF test [3] consisted of two phases; fear acquisition (day 1) and fear extinction (day 2). Fear acquisition was conducted in a Med Associates chamber (30.5 x 24.1 x 21 cm, model VFC-008). Rats were habituated for 10 minutes to the chamber prior to testing. Fear acquisition started with a 2-minute habituation period, followed by five trials (1-minute inter-trial interval) of a 30-second auditory stimulus (85 dB, 2.8 kHz) co-terminated with a 1 second, 0.6 mA foot shock. The shocks were generated using a scrambled shock generator (model ENV-412, Med Associates) connected to the metal grid floor in the chamber. Twenty-four hours later, the fear extinction phase was conducted in a novel, open field context (50 x 50 x 55 cm, clear PMMA) with an external set of speakers that were able to deliver the auditory stimuli (85 dB, 2.8 kHz). The extinction phase started with a 2-minute habituation period, followed by 24 20-second auditory stimuli (85 dB, 2.8 kHz) with a 10 second inter-stimuli interval. Freezing behavior, quantified as the total time (seconds) immobile after auditory stimuli, was manually assessed using BORIS software [4]. The cumulative freezing time for the first and every fourth trial (1st, 4th, 8th, 12th, 16th, 20th and 24$^{th}$) was used to determine SPS-rank for CF.

The PPI test [5, 6] was conducted using a Startle Response System (SR-LAB, SD instruments). Rats were placed in a Perspex tube (8.2 cm diameter, 25 cm length) mounted on a piezoelectric accelerometer. The trial session contained a 5-minute acclimatization phase (70 dB background noise), followed by ten blocks of six stimuli: 120 dB startle stimulus, no stimulus, and four prepulse startle stimuli: 3, 5, 10, and 15 dB above background noise (70 dB)) preceding a 120 dB startle stimulus. Two parameters; the mean P120 maximum acoustic startle (basal startle response), and the average of the PPI at Vmax (PPI Vmax) and across all stimuli (PPI average) were calculated (see equation below). Only the first five blocks were used to prevent habituation effects. All values were corrected using the no stimulus condition to correct for background noise during data acquisition. Only the first five block sessions were used for the ranked selection to prevent habituation, and all values were corrected using the no stimulus condition to correct for background noise during data acquisition.

$$PPI\ Vmax = \frac{\sum_{n \in \{3,5,10,15\}} \left(1 - \left(\frac{\bar{X}_{Vmax(n)} - \bar{X}_{Vmax(no\ stim)}}{\bar{X}_{Vmax(120)} - \bar{X}_{Vmax(no\ stim)}}\right)\right)}{Number\ of\ prepulse\ sessions}$$

$$PPI\ average = \frac{\sum_{n \in \{3,5,10,15\}} (1 - \left(\frac{\bar{X}_{pp(n)} - \bar{X}_{pp(no\ stim)}}{\bar{X}_{pp(120)} - \bar{X}_{pp(no\ stim)}}\right))}{Number\ of\ prepulse\ sessions}$$

***Housing conditions***

With the expected differences in sensitivity to the environment and the known role of environmental conditions on cocaine behavior in rats [7-10], we randomly allocated the high- and low SPS-like rats to three housing conditions; isolated, neutral or enriched. Isolation-housed rats were single-housed in a mouse cage (1284L Eurostandard type II L) without shelter. Neutral-housed rats were pair-housed in a standard rat cage (1290D Eurostandard type III H) with a shelter (Orange rat reteat Bio-Serv™

Thermo Fisher Scientific). Social-housed rats were group-housed (N=6 per cage) in a large cage (75x45x70cm) with a climbing option to a rounded platform (quadrant, radius 38cm, 45 cm high), two large shelters (polyvinyl chloride (PVC), diameter 15 cm), nest-building material (Sizzle nest, Datesand) and plastic toys with bells. The rats were introduced to their new housing conditions six weeks prior to the start of cocaine S/A training.

***Intravenous catheterization***

All high- and low SPS-like rats were implanted with a micro Renathane catheter (Ø0.3 mm ID; Ø0.64 mm OD; #MRE037, Braintree Scientific, Inc.). The catheter was inserted into the right jugular vein under isoflurane anesthesia (5% induction dose, 2-3% maintenance dose) for details: see Verheij et al. 2018 [1]. Analgesic carprofen (10 mg/ml, Rimadyl in non-acidified drinking water) was administered one day prior to the surgery and continued for three days. During the surgery, subcutaneous lidocaine (4 mg/kg, Fresenius Kabi) was injected at the incision sites. Post-surgery, catheters were flushed with 0.2 ml of Saline infused with Cefazoline (175 mg/ml, Fresenius Kabi) and heparin (50 USP, LEO) to prevent infections and blockage.

***Cocaine self-administration***

Cocaine (Bufa Spruyt Hillen, The Netherlands), dissolved in saline (412.5 mg/525 ml of 0.9% saline, Baxter), was administered to freely-moving rats in a standard operant chamber (28 × 26 × 20 cm, Med Associates Inc.) using a syringe pump (Razel Scientific Instruments) and swivel system (for details see: Verheij et al.[11], Caffino et al.[12]). The rats were trained to S/A cocaine (0.5 mg/kg/infusion) at fixed ratio 1 (FR1) for 1h/day for at least 11 days [11, 12]. Cocaine escalation was assessed during 21 days of long access (LgA) cocaine S/A on FR1 for 6h/day [11, 13, 14]. Cocaine motivation was assessed during two progressive ratio (PR) sessions (for details see: Richardson and Roberts[13]). After two weeks of abstinence, cocaine S/A was re-established for 7 days at LgA scheme (under FR1 conditions) before sacrifice and tissue collection. For a complete overview of self-administration session and timing, see fig.2a.

$$Response\ ratio\ of\ PR = \left(5e^{(injection\ number\ x\ 0.2)}\right) - 5$$

***Tissue collection and protein isolation***

Twenty-four hours after the last cocaine S/A reinstatement session, low and high SPS-like rats were sacrificed by decapitation without anesthesia. The brains were collected and immediately frozen on dry ice, and afterwards stored at −80°C. For microdissection of the medial prefrontal cortex (mPFC) subregions the micro-punching technique was used as previously described [15]. In brief, the prelimbic (PLc) and infralimbic (ILc) subregions of the mPFC (coordinates between bregma +4.20 mm and bregma +2.52 mm, Rat Brain Atlas[16]) were punched out from 200 µm slices of the frozen brain using a sterile 1-mm-diameter needle and stored at −80°C. The punches from the right- and left hemispheres of PLc and ILc of a single rat were pooled, homogenized in a cold isotonic buffer [0.32M sucrose buffer pH 7.4; 1 mM HEPES, 0.1 mM PMSF, protease (Roche, Monza, Italy) and phosphatase (Sigma-Aldrich, Milan, Italy) inhibitors] as previously described [17] and sonicated to obtain a whole homogenate extract. Total proteins were measured according to the Bradford Protein Assay procedure (Bio-Rad, Milan, Italy), using bovine serum albumin as calibration standard.

***Western blot analysis***

After protein quantification, the western blots were run as previously described [18]. The proteins (10 µg/sample) were loaded on a sodium dodecyl sulfate-8% (SDS) polyacrylamide gel, electrophoretically separated under reducing conditions, and transferred onto nitrocellulose membranes (GE Healthcare, Milan, Italy). The obtained blots were blocked in I-Block solution (Life Technologies Italia, Italy) in TBS + 0.1% Tween-20 buffer (1 h, room temperature) and incubated with antibodies (see supplementary table 21) against the proteins of interest. Expression levels of every single protein were normalized using its own β-actin loading control, which was detected by evaluating the band density at 43kDa. Optic density (OD) of immunocomplexes was visualized by chemiluminescence using the Chemidoc MP Imaging System (Bio-Rad Laboratories). Gels were run in duplicate and reported values represent the average of two independent runs. We used a correction factor for the second gel (see equation below, with n indicating the number of samples and i indicating each individual control sample) was used for the second gel [15]. The correction factor was determined on the control group, in this case the low SPS-like naïve rats and used for all groups on the second gel.

$$CF_{Gel\ 2} = \frac{1}{n}\sum_{i=1}^{n}\frac{(OD_{Protein\ gel\ 1,i}/OD_{Actin\ gel\ 1,i})}{(OD_{protein\ gel\ 2,i}/OD_{Actin\ gel\ 2,i})}$$

***Statistical analysis of behavioral data***

Behavioral data analysis was performed in IBM SPSS version 30 (IBM software) and RStudio Version 4.4.0 (R Core Team [19]). Behavioral data presented in figures displays mean ± SEM with group sizes reported in the figure text. Group sizes are described in the figure legends. Significance is indicated using (*) for $p < 0.05$, (**) for $p < 0.01$ and (***) for $p < 0.001$.

All SPS selection criteria (EPM, CF, P120s and PPI) were assessed for homoscedasticity using Levene's test (supplementary table 1) and normality using Shapiro-Wilk test (supplementary table 2). To handle heteroscedasticity and non-normality, the EPM, CF, P120s and PPI results were analyzed using a Brown-Forsythe-corrected one-way ANOVA (supplementary table 3), followed by Dunnett T3 post-hoc comparison as recommended for smaller sample sizes [20]. Dimension reduction was conducted using the robust linear discriminant analysis (rLDA) Linda() function [21, 22] from the rrcov package in RStudio as it is more suitable for the assumptions violations mentioned above. The rLDA used prior probabilities of group membership to reflect the observed proportions of rats in each group within the dataset; high SPS-like (0.224), intermediate SPS-like (0.558), and low SPS-like (0.218). Group-specific constants and variable mean per group used in the model are displayed in supplementary table 5. The fractional relative importance was calculated using the within-group variances (supplementary table 6) and the linear discriminant coefficients (supplementary table 7) as described below with *i* referring to a specific variable, *j* referring to all variables, *g* referring to a specific group and *G* referring to all groups:

$$Relative\ Importance_i = (\frac{\sum_g(\sqrt{Variance_i} * coefficent_{i,g}}{G} \ / \ \sum_j \frac{\sum_g(\sqrt{Variance_j} * Coefficent_{i,j})}{G})$$

Cocaine S/A data, both training and LgA, were analyzed in SPSS using linear mixed models (LMMs) due to their ability to handle missing data, higher flexibility and incorporation of random effects and, unlike rmANOVA, do not require sphericity making it suitable for longitudinal data with imbalances [23-26]. Number of cocaine infusions during LgA were normalized across batches based on the number of cocaine infusions received on the final testing day. The LMM included fixed effects for SPS (2 levels), housing condition (3 levels), and time (training: 11 levels, LgA: 21 levels) with a random effect for each individual rat (Subject ID). Time was considered as a repeated measure using covariance type AR(1) heterogeneous and the models were estimated using a Restricted Maximum Likelihood (REML) method. The model specification was as follows: infusions ~ SPS + Condition + Time + SPS*Time + Condition*Time + SPS*Condition + SPS*Condition*Time + (1|Subject). The model fit and corresponding statistical values are displayed in supplementary table 8-10 for cocaine self-administration training and in supplementary table 14-16 for cocaine self-administration long access. Significant fixed effects have been post-hoc analyzed with Bonferroni correction using the estimated marginal means. For cocaine S/A training, post-hoc statistics are displayed for SPS (supplementary figure 11), housing (supplementary figure 12) and SPS*housing interaction (supplementary figure 13). For cocaine S/A LgA, post-hoc statistics are displayed for SPS (supplementary figure 17).

Progressive ratio data was analyzed using two-way ANOVA (between-factor: SPS-level and housing condition; supplementary figure 20) with equality of variances assumed based on Levene's test statistics (supplementary figure 18) and normality assumed based on Shapiro-Wilk test statistics (supplementary figure 19).

***Statistical analysis of molecular data***

Molecular data analysis was performed in GraphPad Prism Version 9.0 (GraphPad Software, Inc.). Subjects were removed from the final dataset if their data deviated from the mean by 2 SDs or more. Group sizes are described in the figure legends. Significance is indicated using (*) for $p < 0.05$, (**) for $p < 0.01$ and (***) for $p < 0.001$.

The molecular changes were analyzed using a two-way (main effects: SPS-like status, cocaine status and interaction SPS*cocaine) using the raw data. ANOVA statistics (supplementary table 22) were adjusted for the number of proteins tested using the Benjamini-Hochberg false discovery rate correction (supplementary table 23). Following FDR correction, relevant main- or interaction effects in the 2x2 design (High OR Low x naive OR cocaine; for clarity see figure 3a) were post-hoc compared using Tukey's multiple comparison test. Data per group is presented as a percentage compared to baseline control (low SPS naïve). Data presented displays the group mean ± S.E.M (figure 3e-f and supplementary figure 3,5). Furthermore, data is presented in a heatmap with a relative comparison between groups displaying the Tukey's post hoc results as mentioned above for high SPS naïve relative to low SPS naïve (figure 3b, baseline differences between high- and low SPS), low SPS cocaine relative to low SPS naïve (figure 3c, cocaine effects within low SPS), and high SPS cocaine relative to high SPS naïve (figure 3d, cocaine effects within high SPS).

### Supplementary Code

#### ***Robust Linear Discriminant Analysis***

```
# Install and load necessary packages if not already installed
if (!require("readxl")) install.packages("readxl")  # For reading Excel files
if (!require("rrcov")) install.packages("rrcov")  # For robust LDA
if (!require("ggplot2")) install.packages("ggplot2")  # For robust LDA
if (!require("writexl")) install.packages("writexl")  # For saving results to Excel
if (!require("caret")) install.packages("caret")  # For confusion matrix and accuracy
if (!require("pheatmap")) install.packages("pheatmap")  # For confusion matrix heatmap
# Load required libraries
library(readxl)
library(rrcov)
library(ggplot2)
library(writexl)
library(caret)  # For confusion matrix and accuracy
library(pheatmap)  # For heatmap of confusion matrix
# Load the data
robust_lda_data <- read_excel("Data.xlsx")  # Adjust the path as necessary
# Fit the Robust LDA model using the Linda function
robust_lda_model_linda <- Linda(SPS ~ PPIgem + Freezing + P120S + `Open arm time`, data = robust_lda_data)
```

```
# Display model summary
summary(robust_lda_model_linda)
# Get the discriminant scores
lda_scores <- predict(robust_lda_model_linda)
# Create a data frame for plotting the LDA results
plot_data <- data.frame(Discriminant1 = lda_scores@x[,1],
                        Discriminant2 = lda_scores@x[,2],
                        SPS = robust_lda_data$SPS)
# Create a scatter plot
ggplot(plot_data, aes(x = Discriminant1, y = Discriminant2, color = SPS)) +
  geom_point(size = 3) +
  labs(title = "Robust LDA: Discriminant Functions",
       x = "Discriminant Function 1",
       y = "Discriminant Function 2") +
  theme_minimal() +
  scale_color_manual(values = c("red", "blue", "green"))
# Save the discriminant coordinates to an Excel file
coordinates_data <- data.frame(Sample = rownames(lda_scores@x),
                               Discriminant1 = lda_scores@x[,1],
                               Discriminant2 = lda_scores@x[,2],
                               SPS = robust_lda_data$SPS)
write_xlsx(coordinates_data, "discriminant_coordinates.xlsx")
# Evaluate the model's predictions against the actual SPS groups
predicted_groups <- predict(robust_lda_model_linda)@classification  # The predicted class labels from the Linda model
actual_groups <- robust_lda_data$SPS  # The actual SPS groups
# Confusion Matrix to compare actual and predicted groups
conf_matrix <- confusionMatrix(predicted_groups, as.factor(actual_groups))
# Display the confusion matrix
print(conf_matrix)
# Visualize the confusion matrix using a heatmap
conf_matrix_table <- as.table(conf_matrix)  # Convert confusion matrix to a table format
pheatmap(conf_matrix_table, display_numbers = TRUE,
```

```
         cluster_rows = FALSE, cluster_cols = FALSE,
         main = "Confusion Matrix Heatmap",
         color = colorRampPalette(c("white", "blue"))(10))
# Accuracy of the model
accuracy <- sum(predicted_groups == actual_groups) / length(actual_groups) * 100
cat("Model Accuracy: ", accuracy, "%\n")

#For variance explained by each LD component
# Extract the covariance matrix from the model
cov_matrix <- robust_lda_model_linda@cov
# Perform Singular Value Decomposition (SVD)
svd_result <- svd(cov_matrix)
# Square the singular values to approximate eigenvalues
eigenvalues <- svd_result$d^2
# Display the eigenvalues
eigenvalues
# Sum of all eigenvalues
total_variance <- sum(eigenvalues)
# Calculate the variance explained by each discriminant function
variance_explained <- (eigenvalues / total_variance) * 100
# Display the results
variance_explained
```

### Supplementary Tables

#### SPS selection criteria statistics

| | Levene Statistics | Df1 | Df2 | Significance |
|---|---|---|---|---|
| **Open arm time (EPM)** | 0.959 | 2 | 162 | 0.385 |
| **Freezing time (CF)** | 4.260 | 2 | 162 | **0.016*** |
| **Startle response (P120s)** | 13.609 | 2 | 162 | **<0.001*** |
| **Prepulse inhibition (PPI)** | 2.857 | 2 | 162 | 0.060 |

**Supplementary table 1** Results of Levene's test for homogeneity of variances assumption for SPS-selection criteria.

| Group | | Statistics | df | significance |
|---|---|---|---|---|
| **EPM** | **Low** | 0.903 | 36 | **0.004*** |
| | **Intermediate** | 0.620 | 92 | **0.548** |
| | **High** | 0.975 | 37 | **<0.001*** |
| **CF** | **Low** | 0.925 | 36 | **0.017*** |
| | **Intermediate** | 0.858 | 92 | **<0.001*** |
| | **High** | 0.548 | 37 | **<0.001*** |
| **P120s** | **Low** | 0.852 | 36 | **<0.001*** |
| | **Intermediate** | 0.909 | 92 | **<0.001*** |
| | **High** | 0.847 | 37 | **<0.001*** |
| **PPI** | **Low** | 0.905 | 36 | **0.005*** |
| | **Intermediate** | 0.743 | 92 | **<0.001*** |
| | **High** | 0.824 | 37 | **<0.001*** |

**Supplementary table 2** Results of Shapiro-Wilk test for normality assumption for SPS-selection criteria.

| | Brown-forsythe F-statistics | Df1 | Df2 | Significance |
|---|---|---|---|---|
| **EPM** | 7.200 | 2 | 158.584 | **0.001*** |
| **CF** | 11.517 | 2 | 86.416 | **<0.001*** |
| **P120S** | 22.039 | 2 | 64.412 | **<0.001*** |
| **PPI** | 7.874 | 2 | 95.651 | **<0.001*** |

**Supplementary table 3** Results of Brown-Forsythe's-corrected one-way ANOVA test for SPS-selection criteria. Brown-Forsythe correction was chosen to deal with violations in homoscedasticity and normality (see supplementary table 2 and 3).

| Group | | Mean difference | Std. error | significance |
|---|---|---|---|---|
| **EPM** | **Low-Intermediate** | 6.33357 | 4.47871 | 0.405 |
| | **Low-High** | 18.11212 | 3.79756 | **<0.001*** |
| | **Intermediate-High** | 11.77855 | 4.16610 | **0.016*** |
| **CF** | **Low-Intermediate** | 22.97186 | 5.28900 | **<0.001*** |
| | **Low-High** | 26.64054 | 6.75637 | **<0.001*** |

| | | | | |
|---|---|---|---|---|
| | **Intermediate-High** | 3.66868 | 4.97845 | 0.843 |
| **P120S** | **Low-Intermediate** | 17.29686 | 5.20724 | **0.004*** |
| | **Low-High** | 61.18258 | 10.11053 | **<0.001*** |
| | **Intermediate-High** | 43.88572 | 10.03395 | **<0.001*** |
| **PPI** | **Low-Intermediate** | 6.81021 | 4.94047 | .431 |
| | **Low-High** | 19.10122 | 4.65519 | **<0.001*** |
| | **Intermediate-High** | 12.29101 | 3.28909 | **<0.001*** |

**Supplementary table 4** Results of Dunnett T3 post-hoc test following Brown-Forsythe's-corrected one-way ANOVA.

### Robust linear discriminant analysis on SPS selection criteria

| | **Group-specific constants** | **Average PPI** | **Freezing time** | **P120s** | **Open arm time** |
|---|---|---|---|---|---|
| **High SPS** | -89.01 | 83.841 | 134.115 | 63.154 | 35.358 |
| **Intermediate SPS** | -66.86 | 72.786 | 122.638 | 40.421 | 40.535 |
| **Low SPS** | -49.08 | 65.018 | 101.242 | 25.930 | 48.387 |

**Supplementary table 5** The group-specific constants and group means per SPS-selection criteria used in the rLDA.

| | **Average PPI** | **Freezing time** | **P120s** | **Open arm time** |
|---|---|---|---|---|
| **Average PPI** | *159.332* | -43.538 | -15.278 | 16.854 |
| **Freezing time** | -43.538 | *262.332* | -54.897 | 10.577 |
| **P120s** | -15.278 | -54.897 | *379.659* | 78.130 |
| **Open arm time** | 16.854 | 10.577 | 78.130 | *225.212* |

**Supplementary table 6** Within-groups covariance matrix used in the rLDA.

| | **Average PPI** | **Freezing time** | **P120s** | **Open arm time** |
|---|---|---|---|---|
| **High SPS** | 0.751 | 0.701 | 0.306 | -0.038 |
| **Intermediate SPS** | 0.644 | 0.618 | 0.216 | 0.027 |
| **Low SPS** | 0.548 | 0.503 | 0.142 | 0.101 |

**Supplementary table 7** Linear discriminant coefficients used in the rLDA.

### Cocaine self-administration training statistics

| | | **Number of levels** | **Number of Parameters** |
|---|---|---|---|
| **Fixed effects** | **Intercept** | 1 | 1 |
| | **SPS** | 2 | 1 |
| | **Housing** | 3 | 2 |
| | **Time** | 11 | 10 |
| | **SPS * Housing** | 6 | 2 |
| | **SPS * Time** | 22 | 10 |
| | **Housing * Time** | 33 | 20 |

| | | | |
|---|---|---|---|
| | **SPS * Housing * Time** | 66 | 20 |
| **Repeated effects** | **Time** | 11 | 12 |
| **Total** | | 155 | 78 |

**Supplementary table 8** Model dimension for linear mixed model of Cocaine S/A training. Covariance structure for repeated measures was heterozygous First-order autoregressive (AR(1)-Heterozygous with random effect of subject ID (N = 35).

| | |
|---|---|
| **-2 Restricted Log Likelihood** | 1735.464224 |
| **Akaike's Information Criterion (AIC)** | 1759.464224 |
| **Schwarz's Bayesian Criterion (BIC)** | 1804.2254198 |

**Supplementary table 9** Information criteria for linear mixed model of Cocaine S/A training.

| Source | Numerator df | Denominator df | F | Sig |
|---|---|---|---|---|
| **Intercept** | 1 | 47.882 | 191.047 | **<0.001*** |
| **SPS** | 1 | 47.882 | 5.466 | **0.024*** |
| **Housing** | 2 | 47.885 | 6.020 | **0.005*** |
| **Time** | 10 | 102.864 | 6.426 | **<0.001*** |
| **SPS * Housing** | 2 | 47.885 | 4.187 | **0.021*** |
| **SPS * Time** | 10 | 102.864 | 2.378 | **0.014*** |
| **Housing * Time** | 20 | 137.479 | 1.556 | 0.073 |
| **SPS * Housing * Time** | 20 | 137.479 | 1.335 | 0.167 |

**Supplementary table 10** Type III test of fixed effects results for linear mixed model of Cocaine S/A training.

| SPS | Mean | Std. error | Mean difference | 95% CI | df | Significance |
|---|---|---|---|---|---|---|
| **High** | 6.607 | 0.586 | 1.912 | 5.429-7.785 | 47.329 | **0.025*** |
| **Low** | 4.695 | 0.571 | -1.912 | 3.548-5.842 | 48.474 | **0.025*** |

**Supplementary table 11** Estimated marginal means and pairwise comparison for SPS (Bonferroni-corrected) for linear mixed model of Cocaine S/A training.

| Housing | Mean | Std. error | Mean difference | 95% CI | df | Significance |
|---|---|---|---|---|---|---|
| **Isolation** | 7.243 | 0.722 | 1.331 (to neutral)<br>3.445 (to social) | 5.7921-<br>8.695 | 47.205 | 0.594 (to neutral)<br>**0.003* (to social)** |
| **Neutral** | 5.912 | 0.704 | -1.331 (to isolation)<br>2.115 (to social) | 4.497-<br>7.327 | 49.114 | 0.594 (to isolation)<br>0.120 (to social) |
| **Social** | 3.798 | 0.698 | -3.445 (to isolation)<br>-2.115 (to neutral) | 2.393-<br>5.202 | 47.376 | **0.003* (to isolation)**<br>0.120 (to neutral) |

**Supplementary table 12** Estimated marginal means and pairwise comparison (Bonferroni-corrected) for housing for linear mixed model of Cocaine S/A training.

| SPS | Housing | Mean | Std. error | 95% CI | df | Significance |
|---|---|---|---|---|---|---|
| **High** | **Isolation** | 9.764 | 1.065 | 7.621-11.906 | 47.023 | **t = 4.20, p=0.0012* (to social)**<br>t = 2.25, p=0.35 (to neutral) |
| | **Neutral** | 6.621 | 0.905 | 4.801-8.440 | 48.186 | t = 2.28, p=0.33 (to social)<br>t = 2.25, p=0.35 (to isolation) |
| | **Social** | 3.436 | 1.065 | 1.294-5.579 | 47.023 | **t = 4.20, p=0.0012* (to isolation)**<br>t = 2.28, p=0.33 (to neutral) |
| **Low** | **Isolation** | 4.723 | 0.974 | 2.764-6.682 | 47.424 | t = 0.42, p=1.00 (to social)<br>t = 0.33, p=1.00 (to neutral) |
| | **Neutral** | 5.204 | 1.079 | 3.036-7.372 | 49.780 | t = 0.74, p=1.00 (to social)<br>t = 0.33, p=1.00 (to isolation) |
| | **Social** | 4.159 | 0.904 | 2.341-5.977 | 47.871 | t = 0.42, p=1.00 (to isolation)<br>t = 0.74, p=1.00 (to isolation) |
| **Isolation** | **High** | 9.764 | 1.065 | 7.621-11.906 | 47.023 | **t = 3.44, p=0.016*** |
| | **Low** | 4.723 | 0.974 | 2.764-6.682 | 47.424 | |
| **Neutral** | **High** | 6.621 | 0.905 | 4.801-8.440 | 48.186 | t = 1.10, p=1.00 |
| | **Low** | 5.204 | 1.079 | 3.036-7.372 | 49.780 | |
| **Social** | **High** | 3.436 | 1.065 | 1.294-5.579 | 47.023 | t = -0.52, p=1.00 |
| | **Low** | 4.159 | 0.904 | 2.341-5.977 | 47.871 | |

**Supplementary table 13** Estimated marginal means and pairwise comparison (Bonferroni-corrected) for SPS*housing interaction for linear mixed model of Cocaine S/A training.

### Cocaine self-administration long access statistics

| | | Number of levels | Number of Parameters |
|---|---|---|---|
| **Fixed effects** | **Intercept** | 1 | 1 |
| | **SPS** | 2 | 1 |
| | **Housing** | 3 | 2 |
| | **Time** | 21 | 20 |
| | **SPS * Housing** | 6 | 2 |
| | **SPS * Time** | 42 | 20 |
| | **Housing * Time** | 63 | 40 |
| | **SPS * Housing * Time** | 126 | 40 |
| **Repeated effects** | **Time** | 21 | 22 |
| **Total** | | 285 | 148 |

**Supplementary table 14** Model dimension for linear mixed model of Cocaine S/A long access. Covariance structure for repeated measures was heterozygous First-order autoregressive (AR(1)-Heterozygous with random effect of subject ID (N = 33).

| | |
|---|---|
| **-2 Restricted Log Likelihood** | 4305.3232871 |
| **Akaike's Information Criterion (AIC)** | 4349.3232871 |
| **Schwarz's Bayesian Criterion (BIC)** | 4444.8111918 |

**Supplementary table 15** Information criteria for linear mixed model of Cocaine S/A long access.

| Source | Numerator df | Denominator df | F | Sig |
|---|---|---|---|---|
| **Intercept** | 1 | 40.808 | 751.236 | **<0.001*** |
| **SPS** | 1 | 40.808 | 9.401 | **0.004*** |
| **Housing** | 2 | 40.808 | 1.585 | 0.217 |
| **Time** | 20 | 192.005 | 4.814 | **<0.001*** |
| **SPS * Housing** | 2 | 40.808 | 0.705 | 0.500 |
| **SPS * Time** | 20 | 192.005 | 1.487 | 0.089 |
| **Housing * Time** | 40 | 263.927 | 1.095 | 0.330 |
| **SPS * Housing * Time** | 40 | 263.927 | 1.250 | 0.156 |

**Supplementary table 16** Type III test of fixed effects results for linear mixed model of Cocaine S/A long access.

| SPS | Mean | Std. error | Mean difference | 95% CI | df | Significance |
|---|---|---|---|---|---|---|
| **High** | 56.862 | 2.708 | 11.442 | 51.393-62.331 | 40.808 | **0.004*** |
| **Low** | 45.420 | 2.568 | -11.442 | 40.233-50.607 | 40.808 | **0.004*** |

**Supplementary table 17** Estimated marginal means and pairwise comparison for SPS (Bonferroni-corrected) for linear mixed model of Cocaine S/A long access.

### Progressive ratio statistics

| Levene's test (progressive ratio) | Df1 | Df2 | Levene's statistics | significance |
|---|---|---|---|---|
| **Progressive ratio (on mean)** | 5 | 26 | 0.420 | 0.830 |

**Supplementary table 18** Levene's test of equality of error variances for progressive ratio test data.

| Group | Statistics | Df | Significance |
|---|---|---|---|
| High SPS – isolation | 0.269 | 5 | 0.537 |
| High SPS – neutral | 0.170 | 6 | 0.473 |
| High SPS – social | 0.212 | 4 | 0.796 |
| Low SPS – isolation | 0.178 | 6 | 0.909 |
| Low SPS – neutral | 0.314 | 5 | 0.122 |
| Low SPS – social | 0.265 | 6 | 0.333 |
| High SPS – all housing | 0.113 | 15 | 0.987 |

| | | | |
|---|---|---|---|
| Low SPS – all housing | 0.098 | 17 | 0.612 |

**Supplementary table 19** Results of Shapiro-Wilk test for normality assumption for progressive ratio test data for all groups (TOP: SPS*housing) and significant main effects from univariate ANOVA (Bottom: SPS).

| **Effect** | **F-statistics** | **Df1** | **Df2** | **Significance** |
|---|---|---|---|---|
| SPS | 6.205 | 1 | 26 | **0.019*** |
| Housing | 1.051 | 2 | 26 | 0.364 |
| SPS*Housing | 0.088 | 2 | 26 | 0.916 |

**Supplementary table 20** Results of two-way ANOVA for progressive ratio data with SPS- and housing effects and interaction

### Western blot antibody table

| **Antibody name** | **Dilution** | **Company** | **RRID** |
|---|---|---|---|
| anti-vGlut1 | 1:1000 | Cell Signalling Technology Inc. | AB_2797887 |
| anti-vGAT | 1:1000 | GeneTex International Corporation | AB_10619521 |
| anti-GLT-1 | 1:5000 | AbCam | AB_1566262 |
| anti-GAD67 | 1:2000 | AbCam | AB_448990 |
| anti-GAT3 | 1:2000 | AbCam | AB_11127719 |
| anti-GluA1 | 1:2000 | Cell Signaling Technology Inc. | AB_2732897 |
| anti-GluA2 | 1:2000 | Cell Signaling Technology Inc. | AB_2650557 |
| anti-GluN1 | 1:1000 | Cell Signaling Technology Inc. | AB_1904067 |
| anti-GluN2A | 1:1000 | Cell Signaling Technology Inc. | AB_2112295 |
| anti-GluN2B | 1:1000 | Cell Signaling Technology Inc. | AB_2798506 |
| anti-SAP102 | 1:1000 | Cell Signaling Technology Inc. | AB_2799325 |
| anti-GRIP | 1:1000 | AbCam | AB_2091910 |
| anti-PSD95 | 1:1000 | Cell Signaling Technology Inc. | AB_2292883 |
| anti-Gephyrin | 1:1000 | Synaptic System | AB_2619837 |
| anti-Neuroligin-1 | 1:1000 | Synaptic System | AB_887746 |
| anti-Neuroligin-2 | 1:1000 | Synaptic System | AB_2619813 |
| anti-β-Actin | 1:10000 | Sigma-Aldrich | AB_476697 |

**Supplementary table 21** List of antibodies used in western blot analysis of prelimbic- and infralimbic cortex with dilution, company and research resource identifier (RRID).

| *P*rotein | ILc interaction | ILc SPS | ILc Cocaine | PLc interaction | PLc SPS | PLc Cocaine |
|---|---|---|---|---|---|---|
| **vGluT1** | $F_{1, 18} = 0.008$, $p = 0.926$ | $F_{1, 18} = 0.006$, $p = 0.936$ | $F_{1, 18} = 0.300$, $p = 0.5906$ | **$F_{1, 18} = 7.096$, $p = 0.0158$** | $F_{1, 18} = 0.677$, $p = 0.4214$ | **$F_{1, 18} = 4.944$, $p = 0.0392$** |
| **vGAT** | **$F_{1, 17} = 14.91$, $p = 0.0013$** | **$F_{1, 17} = 6.523$, $p = 0.0205$** | $F_{1, 17} = 2.563$, $p = 0.1278$ | **$F_{1, 18} = 10.84$, $p = 0.0041$** | **$F_{1, 18} = 6.004$, $p = 0.0247$** | **$F_{1, 18} = 8{,}064$, $p = 0.0109$** |
| **vGlut /vGAT** | **$F_{1, 17} = 12.90$, $p = 0.0023$** | **$F_{1, 17} = 6.531$, $p = 0.0205$** | $F_{1, 17} = 2.804$, $p = 0.1123$ | $F_{1, 18} = 0.749$, $p = 0.3982$ | **$F_{1, 18} = 9.666$, $p = 0.0061$** | **$F_{1, 18} = 19.28$, $p = 0.0004$** |
| **GLT1** | **$F_{1, 18} = 21.47$, $p = 0.0002$** | $F_{1, 18} = 0.391$, $p = 0.5397$ | **$F_{1, 18} = 5.944$, $p = 0.0254$** | $F_{1, 18} = 0.104$, $p = 0.7505$ | $F_{1, 18} = 2.385$, $p = 0.1399$ | $F_{1, 18} = 0.268$, $p = 0.6112$ |
| **GAD67** | **$F_{1, 18} = 9.118$, $p = 0.0074$** | **$F_{1, 18} = 25.72$, $p < 0.0001$** | **$F_{1, 18} = 17.97$, $p = 0.0005$** | **$F_{1, 18} = 19.95$, $p = 0.0003$** | $F_{1, 18} = 2.209$, $p = 0.1545$ | **$F_{1, 18} = 0.011$, $p = 0.9187$** |
| **GAT3** | **$F_{1, 18} = 30.32$, $p<0.0001$** | **$F_{1, 18} = 11.67$, $p = 0.0031$** | $F_{1, 18} = 1.761$, $p = 0.2011$ | **$F_{1, 18} = 11.87$, $p = 0.0029$** | **$F_{1, 18} = 22.38$, $p = 0.0002$** | $F_{1, 18} = 0.367$, $p = 0.5524$ |
| **GluA1** | **$F_{1, 18} = 48.18$, $p < 0.0001$** | $F_{1, 18} = 2.138$, $p = 0.1609$ | **$F_{1, 18} = 5.142$, $p = 0.0359$** | **$F_{1, 18} = 9.471$, $p = 0.0065$** | **$F_{1, 18} = 4.715$, $p = 0.0435$** | **$F_{1, 18} = 4.112$, $p = 0.0577$** |
| **GluA2** | **$F_{1, 18} = 7.971$, $p = 0.0113$** | **$F_{1, 18} = 10.12$, $p = 0.0052$** | **$F_{1, 18} = 11.52$, $p = 0.0032$** | **$F_{1, 18} = 22.42$, $p = 0.0002$** | **$F_{1, 18} = 5.063$, $p = 0.0372$** | **$F_{1, 18} = 10.42$, $p = 0.0047$** |
| **GluA1 /GluA2** | **$F_{1, 18} = 26.66$, $p < 0.0001$** | $F_{1, 18} = 1.569$, $p = 0.2264$ | $F_{1, 18} = 0.302$, $p = 0.5895$ | **$F_{1, 18} = 5.347$, $p = 0.0328$** | **$F_{1, 18} = 10.60$, $p = 0.0044$** | **$F_{1, 18} = 20.94$, $p = 0.0002$** |
| **GluN1** | **$F_{1, 18} = 24.67$, $p < 0.0001$** | $F_{1, 18} = 0.568$, $p = 0.4607$ | $F_{1, 18} = 0.106$, $p = 0.7483$ | **$F_{1, 18} = 15.79$, $p = 0.0009$** | $F_{1, 18} = 0.181$, $p = 0.6760$ | $F_{1, 18} = 2.721$, $p = 0.1164$ |
| **GluN2A** | **$F_{1, 18} = 4.583$, $p = 0.0462$** | **$F_{1, 18} = 4.855$, $p = 0.0408$** | **$F_{1, 18} = 10.67$, $p = 0.0043$** | $F_{1, 18} = 1.093$, $p = 0.3096$ | **$F_{1, 18} = 5.276$, $p = 0.0338$** | **$F_{1, 18} = 14.06$, $p = 0.0015$** |
| **GluN2B** | **$F_{1, 18} = 44.12$, $p < 0.0001$** | $F_{1, 18} = 0.021$, $p = 0.8869$ | $F_{1, 18} = 0.158$, $p = 0.6954$ | **$F_{1, 18} = 29.88$, $p < 0.0001$** | $F_{1, 18} = 2.356$, $p = 0.1422$ | $F_{1, 18} = 2.439$, $p = 0.1358$ |
| **GluN2A /GluN2B** | **$F_{1, 18} = 5.056$, $p = 0.0373$** | $F_{1, 18} = 3.189$, $p = 0.0910$ | **$F_{1, 18} = 5.969$, $p = 0.0251$** | **$F_{1, 18} = 19.29$, $p = 0.0004$** | $F_{1, 18} = 2.317$, $p = 0.1454$ | **$F_{1, 18} = 20.15$, $p = 0.0003$** |
| **SAP102** | **$F_{1, 18} = 14.15$, $p = 0.0014$** | $F_{1, 18} = 0.044$, $p = 0.8358$ | $F_{1, 18} = 0.459$, $p = 0.5066$ | **$F_{1, 18} = 17.62$, $p = 0.0005$** | $F_{1, 18} = 0.151$, $p = 0.7022$ | **$F_{1, 18} = 16.03$, $p = 0.0008$** |
| **GRIP** | $F_{1, 18} = 0.276$, $p = 0.6059$ | $F_{1, 18} = 1{,}781$, $p = 0.1986$ | $F_{1, 18} = 0.130$, $p = 0.7223$ | **$F_{1, 18} = 10.46$, $p = 0.0046$** | **$F_{1, 18} = 29.29$, $p < 0.0001$** | **$F_{1, 18} = 29.44$, $p < 0.0001$** |
| **PSD95** | **$F_{1, 18} = 9.423$, $p = 0.0066$** | **$F_{1, 18} = 7.740$, $p = 0.0123$** | **$F_{1, 18} = 16.18$, $p = 0.0008$** | **$F_{1, 18} = 6.279$, $p = 0.0220$** | **$F_{1, 18} = 21.25$, $p = 0.0002$** | **$F_{1, 18} = 11.02$, $p = 0.0038$** |
| **Gephyrin** | **$F_{1, 18} = 4.622$, $p = 0.0454$** | $F_{1, 18} = 0.003$, $p = 0.9581$ | $F_{1, 18} = 3.141$, $p = 0.0933$ | $F_{1, 18} = 1.046$, $p = 0.3200$ | $F_{1, 18} = 0.005$, $p = 0.9439$ | $F_{1, 18} = 0.775$, $p = 0.3904$ |
| **NLGN1** | **$F_{1, 18} = 4.698$, $p = 0.0438$** | $F_{1, 18} = 0.650$, $p = 0.4305$ | **$F_{1, 18} = 6.137$, $p = 0.0234$** | $F_{1, 18} = 0.024$, $p = 0.8780$ | $F_{1, 18} = 1.539$, $p = 0.2306$ | **$F_{1, 18} = 20.64$, $p = 0.0003$** |
| **NLGN2** | **$F_{1, 18} = 9.867$, $p = 0.0056$** | $F_{1, 18} = 1.719$, $p = 0.2063$ | **$F_{1, 18} = 5.876$, $p = 0.0261$** | **$F_{1, 18} = 9.654$, $p = 0.0061$** | $F_{1, 18} = 0.030$, $p = 0.8653$ | **$F_{1, 18} = 9.404$, $p = 0.0066$** |

**Supplementary table 22** F-value and corresponding *p*-value of interaction (SPS*Cocaine) and main effects (SPS or Cocaine) of the two-way ANOVA western blot protein data. Significant results highlighted in bold.

| *P*rotein | ILc interaction | ILc SPS | ILc Cocaine | PLc interaction | PLc SPS | PLc Cocaine |
|---|---|---|---|---|---|---|
| **vGluT1** | $p$ = 0.926<br>$R_{19}$, α=0.05 | $p$ = 0.936<br>$R_{18}$, α=0.047 | $p$ = 0.5906<br>$R_{16}$, α=0.042 | **$p$ = 0.0158<br>$R_{12}$, α=0.032** | $p$ = 0.4214<br>$R_{15}$, α=0.039 | $p$ = 0.0392<br>$R_{12}$, α=0.032 |
| **vGAT** | **$p$ = 0.0013<br>$R_7$, α=0.018** | $p$ = 0.0205<br>$R_5$, α=0.013 | $p$ = 0.1278<br>$R_{12}$, α=0.032 | **$p$ = 0.0041<br>$R_8$, α=0.021** | $p$ = 0.0247<br>$R_6$, α=0.016 | **$p$ = 0.0109<br>$R_{11}$, α=0.029** |
| **vGlut /vGAT** | **$p$ = 0.0023<br>$R_9$, α=0.024** | $p$ = 0.0205<br>$R_6$, α=0.016 | $p$ = 0.1123<br>$R_{11}$, α=0.029 | $p$ = 0.3982<br>$R_{17}$, α=0.045 | **$p$ = 0.0061<br>$R_5$, α=0.013** | **$p$ = 0.0004<br>$R_5$, α=0.013** |
| **GLT1** | **$p$ = 0.0002<br>$R_6$, α=0.016** | $p$ = 0.5397<br>$R_{15}$, α=0.039 | $p$ = 0.0254<br>$R_7$, α=0.018 | $p$ = 0.7505<br>$R_{18}$, α=0.047 | $p$ = 0.1399<br>$R_{10}$, α=0.026 | $p$ = 0.6112<br>$R_{18}$, α=0.047 |
| **GAD67** | **$p$ = 0.0074<br>$R_{12}$, α=0.032** | **$p$ < 0.0001<br>$R_1$, α=0.003** | **$p$ = 0.0005<br>$R_1$, α=0.003** | **$p$ = 0.0003<br>$R_3$, α=0.008** | $p$ = 0.1545<br>$R_{13}$, α=0.034 | $p$ = 0.9187<br>$R_{19}$, α=0.05 |
| **GAT3** | **$p$ < 0.0001<br>$R_1$, α=0.003** | **$p$ = 0.0031<br>$R_2$, α=0.005** | $p$ = 0.2011<br>$R_{13}$, α=0.034 | **$p$ = 0.0029<br>$R_7$, α=0.018** | **$p$ = 0.0002<br>$R_2$, α=0.005** | $p$ = 0.5524<br>$R_{17}$, α=0.045 |
| **GluA1** | **$p$ < 0.0001<br>$R_2$, α=0.005** | $p$ = 0.1609<br>$R_9$, α=0.024 | $p$ = 0.0359<br>$R_9$, α=0.024 | **$p$ = 0.0065<br>$R_{11}$, α=0.029** | $p$ = 0.0435<br>$R_9$, α=0.024 | $p$ = 0.0577<br>$R_{13}$, α=0.034 |
| **GluA2** | **$p$ = 0.0113<br>$R_{13}$, α=0.034** | **$p$ = 0.0052<br>$R_3$, α=0.008** | **$p$ = 0.0032<br>$R_3$, α=0.008** | **$p$ = 0.0002<br>$R_2$, α=0.005** | $p$ = 0.0372<br>$R_8$, α=0.021 | **$p$ = 0.0047<br>$R_9$, α=0.024** |
| **GluA1 /GluA2** | **$p$ < 0.0001<br>$R_3$, α=0.008** | $p$ = 0.2264<br>$R_{12}$, α=0.032 | $p$ = 0.5895<br>$R_{15}$, α=0.039 | **$p$ = 0.0328<br>$R_{14}$, α=0.037** | **$p$ = 0.0044<br>$R_4$, α=0.011** | **$p$ = 0.0002<br>$R_2$, α=0.005** |
| **GluN1** | **$p$ < 0.0001<br>$R_4$, α=0.011** | $p$ = 0.4607<br>$R_{14}$, α=0.037 | $p$ = 0.7483<br>$R_{19}$, α=0.05 | **$p$ = 0.0009<br>$R_6$, α=0.016** | $p$ = 0.6760<br>$R_{16}$, α=0.042 | $p$ = 0.1164<br>$R_{14}$, α=0.037 |
| **GluN2A** | $p$ = 0.0462<br>$R_{17}$, α=0.045 | $p$ = 0.0408<br>$R_7$, α=0.018 | **$p$ = 0.0043<br>$R_4$, α=0.011** | $p$ = 0.3096<br>$R_{15}$, α=0.039 | $p$ = 0.0338<br>$R_7$, α=0.018 | **$p$ = 0.0015<br>$R_7$, α=0.018** |
| **GluN2B** | **$p$ < 0.0001<br>$R_5$, α=0.013** | $p$ = 0.8869<br>$R_{17}$, α=0.045 | $p$ = 0.6954<br>$R_{17}$, α=0.045 | **$p$ < 0.0001<br>$R_1$, α=0.003** | $p$ = 0.1422<br>$R_{11}$, α=0.029 | $p$ = 0.1358<br>$R_{15}$, α=0.039 |
| **GluN2A /GluN2B** | $p$ = 0.0373<br>$R_{14}$, α=0.037 | $p$ = 0.0910<br>$R_8$, α=0.021 | $p$ = 0.0251<br>$R_6$, α=0.016 | **$p$ = 0.0004<br>$R_4$, α=0.011** | $p$ = 0.1454<br>$R_{12}$, α=0.032 | **$p$ = 0.0003<br>$R_3$, α=0.008** |
| **SAP102** | **$p$ = 0.0014<br>$R_8$, α=0.021** | $p$ = 0.8358<br>$R_{16}$, α=0.042 | $p$ = 0.5066<br>$R_{14}$, α=0.037 | **$p$ = 0.0005<br>$R_5$, α=0.013** | $p$ = 0.7022<br>$R_{17}$, α=0.045 | **$p$ = 0.0008<br>$R_6$, α=0.016** |
| **GRIP** | $p$ = 0.6059<br>$R_{18}$, α=0.047 | $p$ = 0.1986<br>$R_{10}$, α=0.026 | $p$ = 0.7223<br>$R_{18}$, α=0.047 | **$p$ = 0.0046<br>$R_9$, α=0.024** | **$p$ < 0.0001<br>$R_1$, α=0.003** | **$p$ < 0.0001<br>$R_1$, α=0.003** |
| **PSD95** | **$p$ = 0.0066<br>$R_{11}$, α=0.029** | $p$ = 0.0123<br>$R_4$, α=0.011 | **$p$ = 0.0008<br>$R_2$, α=0.005** | **$p$ = 0.0220<br>$R_{13}$, α=0.034** | **$p$ = 0.0002<br>$R_3$, α=0.008** | **$p$ = 0.0038<br>$R_8$, α=0.021** |
| **Gephyrin** | $p$ = 0.0454<br>$R_{16}$, α=0.042 | $p$ = 0.9581<br>$R_{19}$, α=0.05 | $p$ = 0.0933<br>$R_{10}$, α=0.026 | $p$ = 0.3200<br>$R_{16}$, α=0.042 | $p$ = 0.9439<br>$R_{19}$, α=0.05 | $p$ = 0.3904<br>$R_{16}$, α=0.042 |
| **NLGN1** | $p$ = 0.0438<br>$R_{15}$, α=0.039 | $p$ = 0.4305<br>$R_{13}$, α=0.034 | $p$ = 0.0234<br>$R_5$, α=0.013 | $p$ = 0.8780<br>$R_{19}$, α=0.05 | $p$ = 0.2306<br>$R_{14}$, α=0.037 | **$p$ = 0.0003<br>$R_4$, α=0.011** |
| **NLGN2** | **$p$ = 0.0056<br>$R_{10}$, α=0.026** | $p$ = 0.2063<br>$R_{11}$, α=0.029 | $p$ = 0.0261<br>$R_8$, α=0.021 | **$p$ = 0.0061<br>$R_{10}$, α=0.026** | $p$ = 0.8653<br>$R_{18}$, α=0.047 | **$p$ = 0.0066<br>$R_{10}$, α=0.026** |

**Supplementary table 23** Benjamini-Hochberg correction factor applied to control false discovery rate (α=0.0X) for the 19 different comparisons performed on the western blot protein data. *p*-values derived from the two-way ANOVA have been corrected based on their respective rank ($R_x$) within the interaction or main ($R_1$ till $R_{19}$)

with the alpha level adjusted ($\alpha=0.05/(p\text{-value Rank}/N_{comparisons})$) with $N_{comparisons} = 19$. Significant results are highlighted in bold.

**Supplementary Figures**

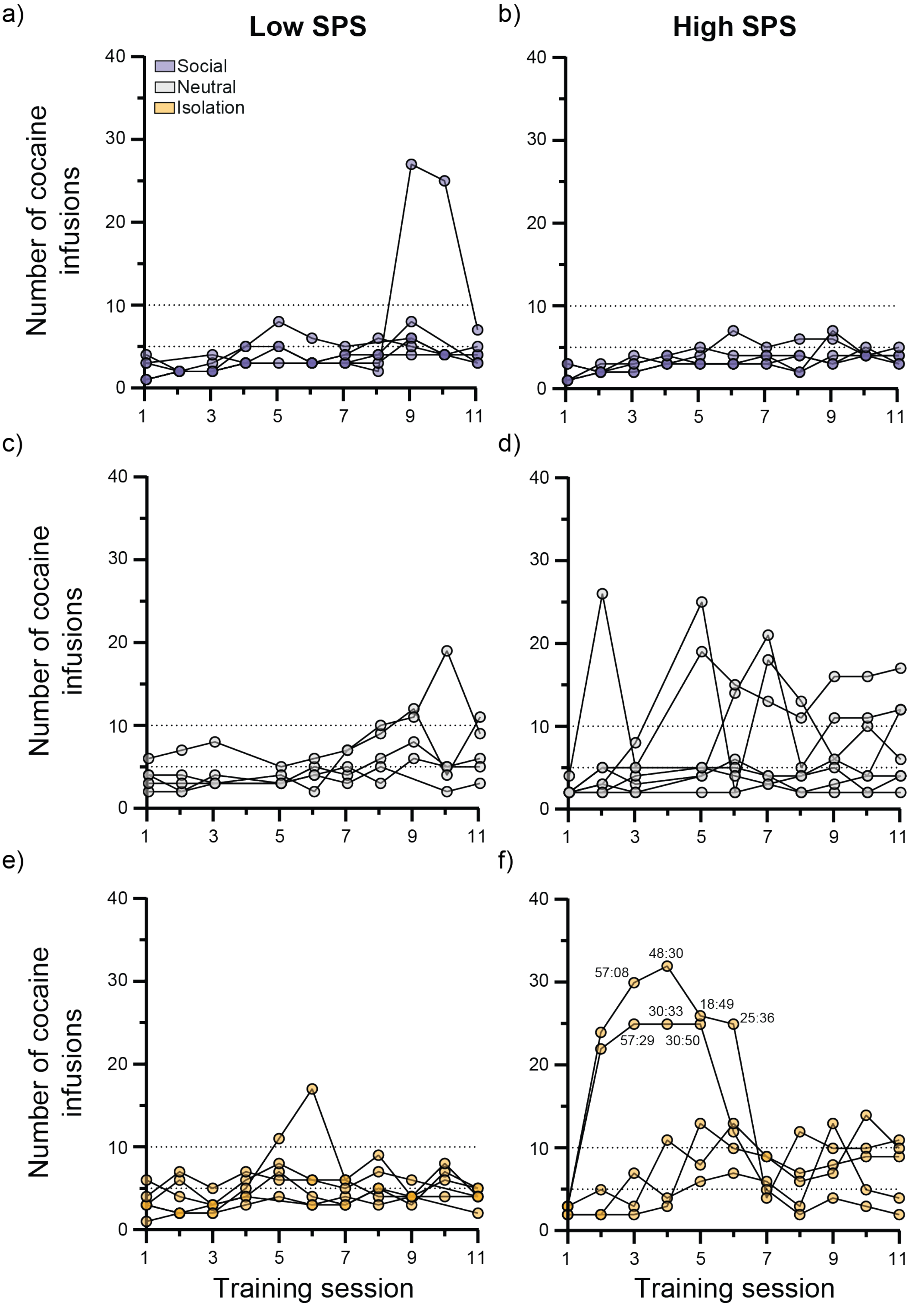

**Supplementary figure 1** Number of self-administered cocaine infusions during 11 days of self-administration training with left panels (a, c, e) representing low SPS-like- and right panels (b, d, f) representing high SPS-like rats. Horizontally, top panels (a, b) represent social housing, middle panels (c, d) represent neutral housing and bottom panels (e, f) represent isolation housing. Each line represents a single rat. Time in high SPS-isolation graph (bottom, right) indicates end time (minute: second) when training was stopped to prevent too much cocaine intake during training. $N_{\text{low SPS - social}} = 6$, $N_{\text{low SPS - neutral}} = 5$, $N_{\text{low SPS - isolation}} = 6$, $N_{\text{high SPS - social}} = 5$, $N_{\text{high SPS - neutral}} = 7$, $N_{\text{high SPS - isolation}} = 5$. Datapoints have transparent color coding to allow visualization of overlapping datapoints.

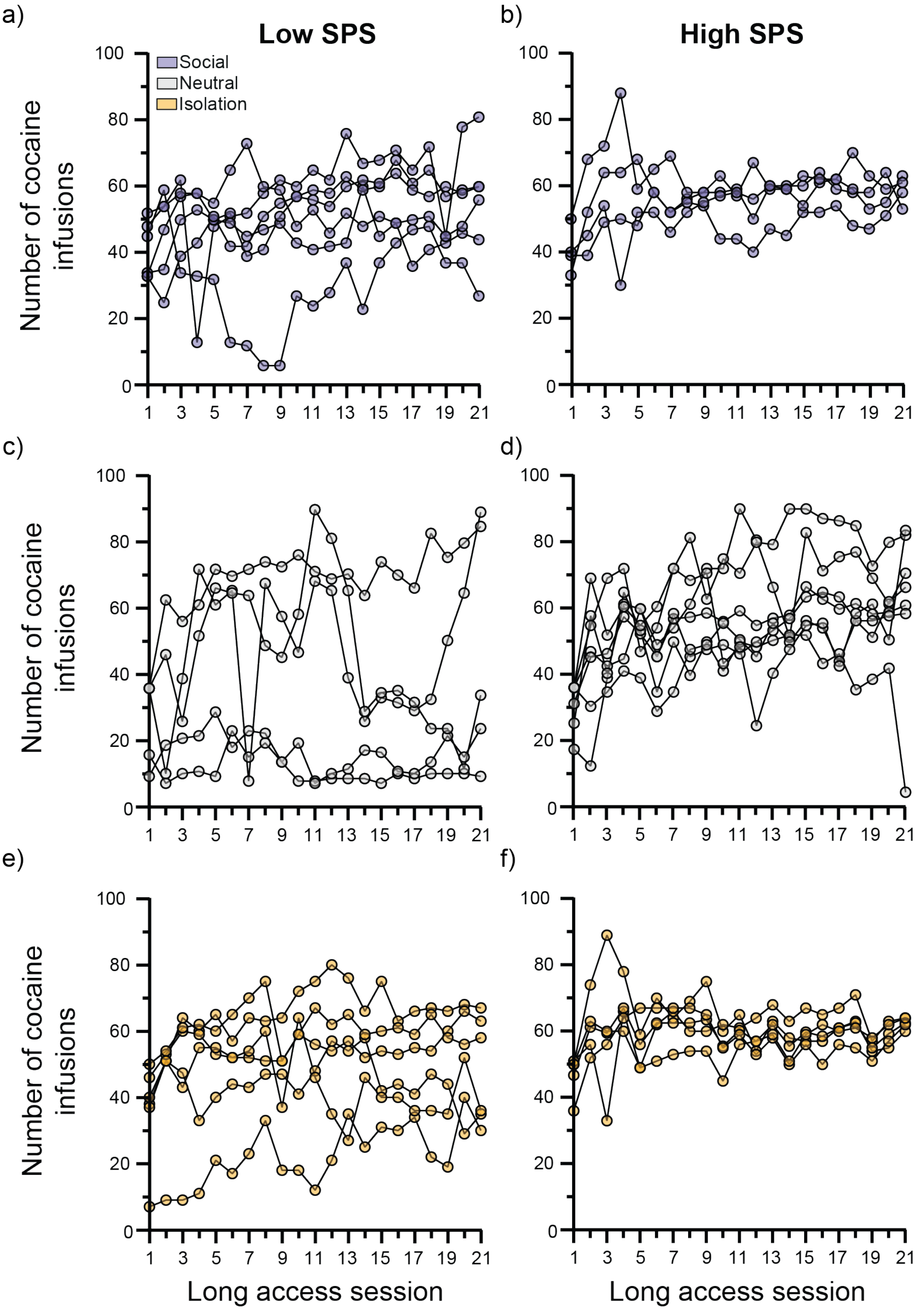

**Supplementary figure 2** Number of self-administered cocaine infusions during 21 days of Long-access self-administration with left panels (a, c, e) representing low SPS-like- and right panels (b, d, f) representing high SPS-like rats. Horizontally, top panels (a, b) represent social housing, middle panels (c, d) represent neutral housing and bottom panels (e, f) represent isolation housing. Each line represents a single rat. $N_{\text{low SPS - social}} = 6$, $N_{\text{low SPS -}}$

$_{\text{neutral}}$ = 5, $N_{\text{low SPS - isolation}}$ = 6, $N_{\text{high SPS - social}}$ = 4, $N_{\text{high SPS - neutral}}$ = 7, $N_{\text{high SPS - isolation}}$ = 5. Datapoints have transparent color coding to allow visualization of overlapping datapoints.

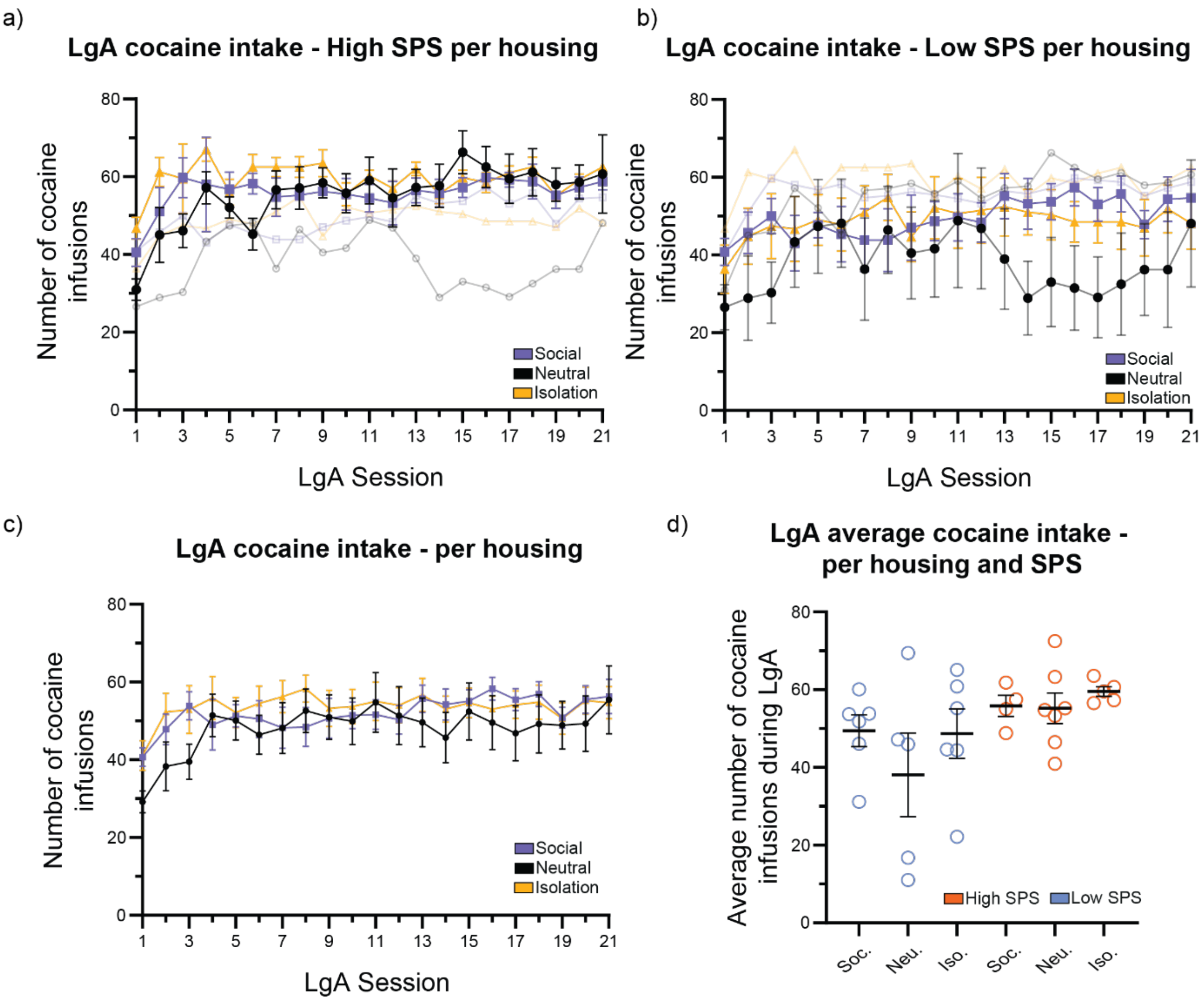

**Supplementary figure 3** a) Number of self-administered cocaine infusions during 21 days of LgA per housing condition and SPS with in focus (full color) high SPS and out focus for comparison (transparent) low SPS. b) Number of self-administered cocaine infusions during 21 days of LgA per housing condition and SPS with in focus (full color) low SPS and out focus for comparison (transparent) high SPS. c) Number of self-administered cocaine infusions during 21 days of LgA per housing condition. d) Average number of cocaine infusions during LgA per housing condition and SPS. The number of rats per group: $N_{\text{low SPS - social}}$ = 6, $N_{\text{low SPS - neutral}}$ = 5, $N_{\text{low SPS - isolation}}$ = 6, $N_{\text{high SPS - social}}$ = 4, $N_{\text{high SPS - neutral}}$ = 7, $N_{\text{high SPS - isolation}}$ = 5. Group means ± SEM are shown.

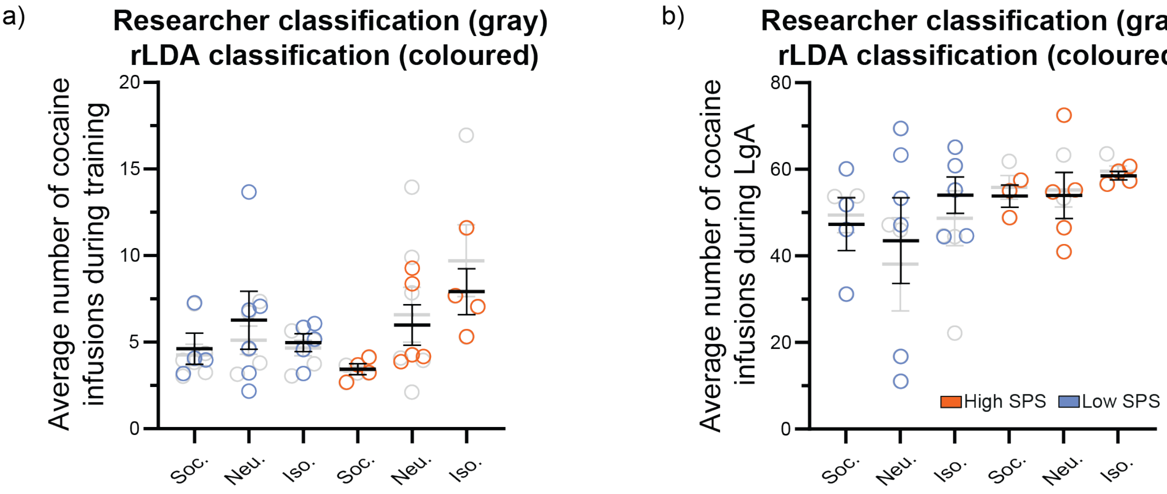

**Supplementary figure 4** Average number of cocaine infusions during training (a) and long access (b) when selecting the cases according to the robust linear discriminant analysis. In focus (full color) the new distribution when considering the rLDA selection (see figure 1-h). In gray, the researcher-based selection (25% weight per selection criteria test, fig1). Due to only researcher assigned high and low SPS-like rats being tested in cocaine S/A paradigm, this would result in a loss of data and thereby statistical power.

Overall, this reclassification resulted in a loss of 8 out of 35 (23%) datapoints for self-administration (S/A) training and 7 out of 33 (21%) datapoints for cocaine LgA, thereby reducing statistical power. When groups were redefined using the rLDA classification, the pattern of estimated marginal means remained in the same direction and of comparable magnitude. Although standard errors did not overlap between groups, the wider 95% confidence intervals overlapped, indicating increased uncertainty rather than a reversal of the effects.

The SPS effect during S/A LgA was no longer statistically significant ($p = 0.132$), but the differences between groups (Mean ± SE [95% CI]: High SPS = 55.4 ± 3.5 [48.4–62.5]; Low SPS = 48.3 ± 3.1 [42.0–54.5]) indicate a similar pattern. Similarly, the SPS × Housing interaction was no longer significant ($p = 0.164$), but the differences per comparison (Mean ± SE [95% CI]: High SPS–isolation = 8.0 ± 1.2 [5.6–10.3] vs. Low SPS–isolation = 4.9 ± 1.0 [2.8–7.0] OR High SPS–social = 3.4 ± 1.2 [1.1–5.8]) were in the same direction, consistent with the pattern observed in the main analysis.

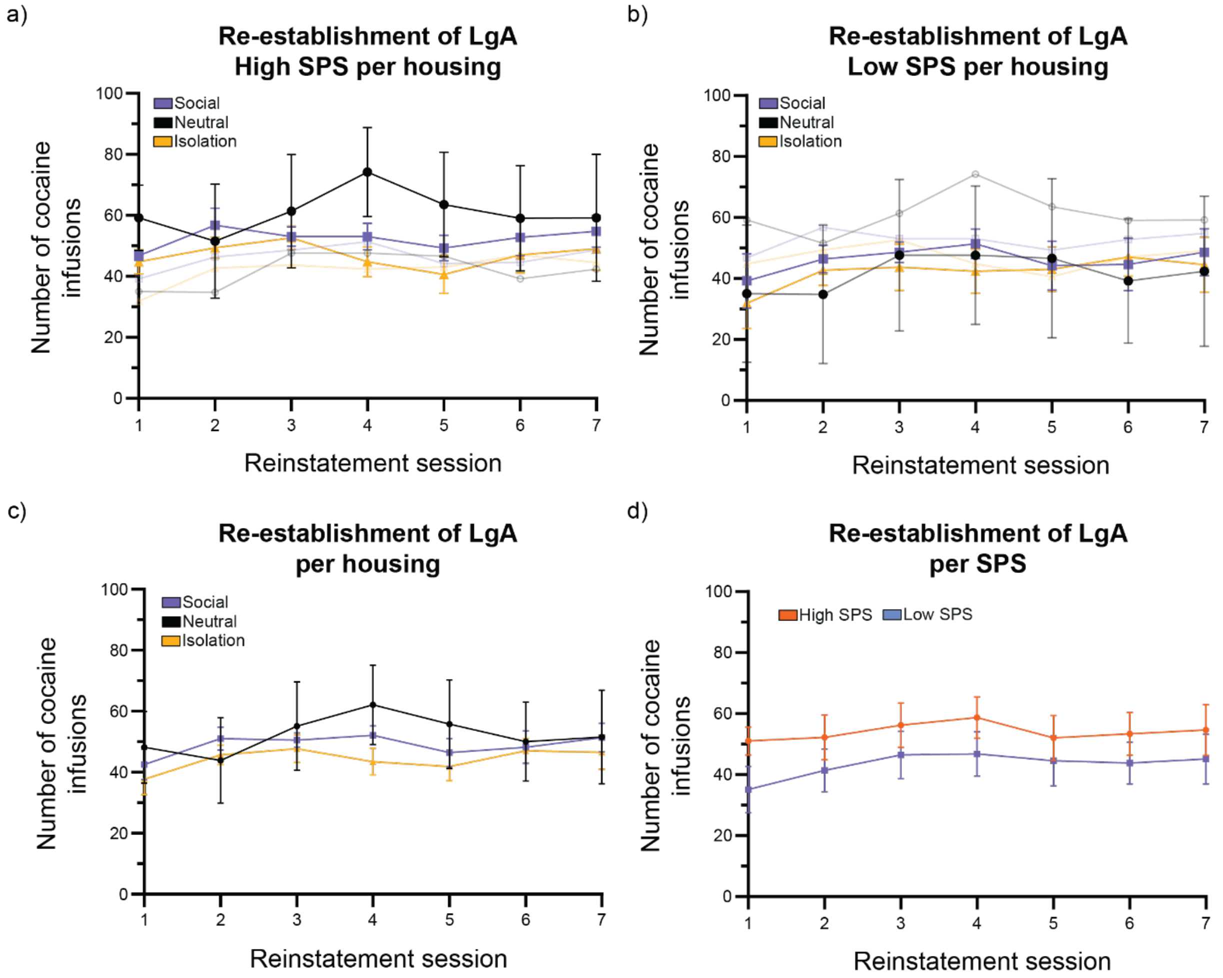

**Supplementary figure 5** a, b) Number of self-administered cocaine infusions during 7 days of LgA re-establishment per housing condition for high SPS-like (a) and low SPS-like (b) rats. In transparent similar color coding, the other group is represented. c) number of self-administered cocaine infusions during 7 days of reinstatement per housing condition (both high- and low SPS-like groups combined). d) The number of cocaine infusions during 7 days of reinstatement per SPS (all housing conditions combined). Linear mixed model revealed a significant main effect of time and significant interaction of time * housing condition, independent of SPS effects. SPS was no longer found to be a significant effect (p=0.344) with high overlap between groups (Mean ± SE [95% CI]: High SPS = 53.4 ± 7.5 [37.9–68.9]; Low SPS = 43.4 ± 7.2 [28.5–58.2]). The number of rats per group: $N_{low\ SPS\ -\ social}$ = 5, $N_{low\ SPS\ -\ neutral}$ = 5, $N_{low\ SPS\ -\ isolation}$ = 6, $N_{high\ SPS\ -\ social}$ = 4, $N_{high\ SPS\ -\ neutral}$ = 6, $N_{high\ SPS\ -\ isolation}$ = 5. Group means ± SEM are shown.

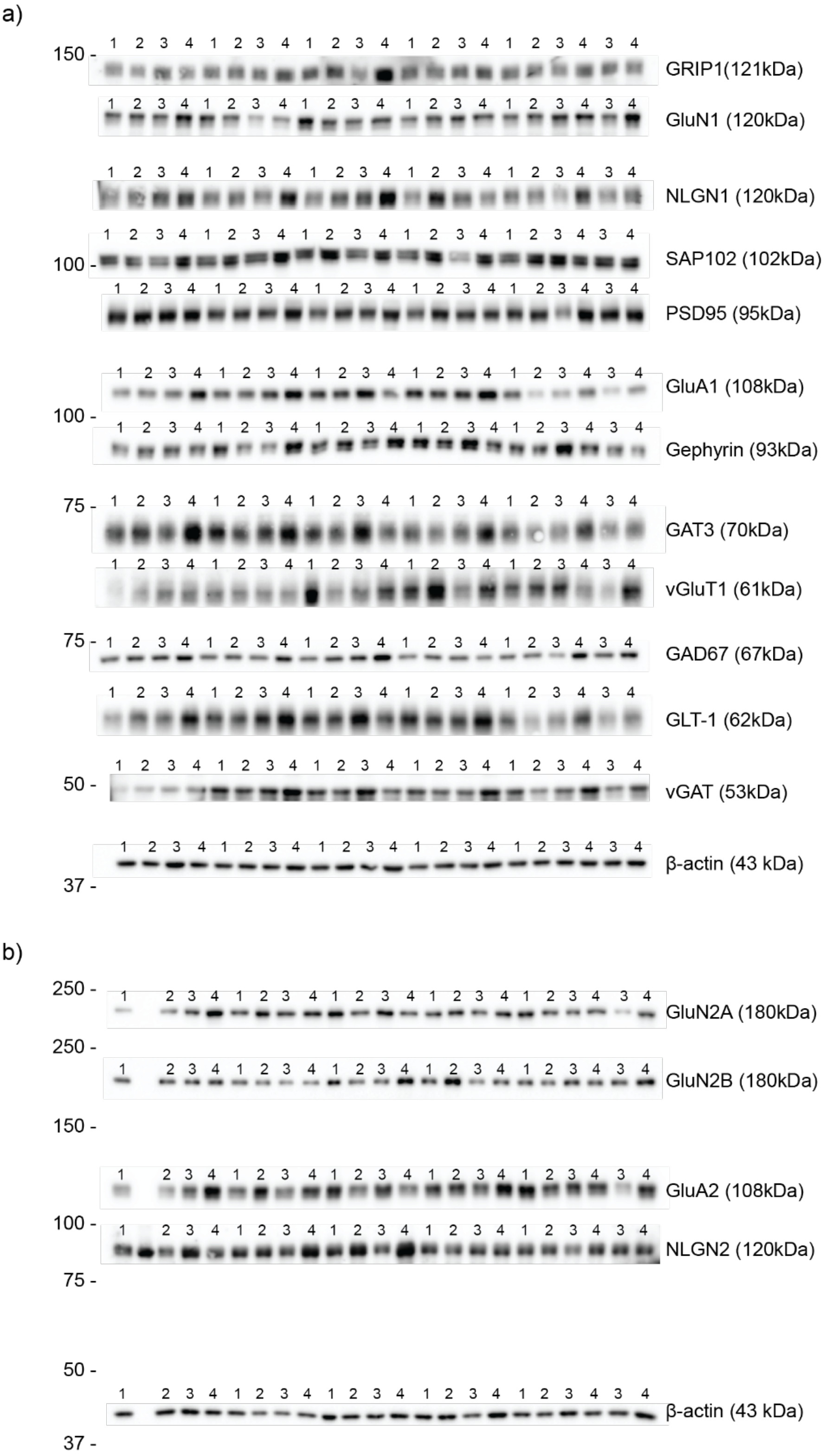

**Supplementary figure 6** Cropped immunoblot related to the protein expression levels of (a) GAD67 (67 kDa), GAT3 (70 kDa), Gephyrin (93 kDa), GLT-1 (62 kDa), GluA1 (108 kDa), GluN1 (120 kDa), GRIP (121 kDa), NLGN1 (120 kDa), PSD95 (95 kDa), SAP102 (102 kDa), VGAT (53 kDa), VGlut1 (61 kDa) and β-actin (43 kDa), and (b) GluN2A (180 kDa), GluN2B (180 kDa), GluA2 (108 kDa), NLGN2 (120 kDa) and β-actin (43 kDa) (b) measured in the homogenate of infralimbic cortex of Low SPS naive (1), Low SPS cocaine (2), High SPS naive (3) and High SPS cocaine (4) rats.

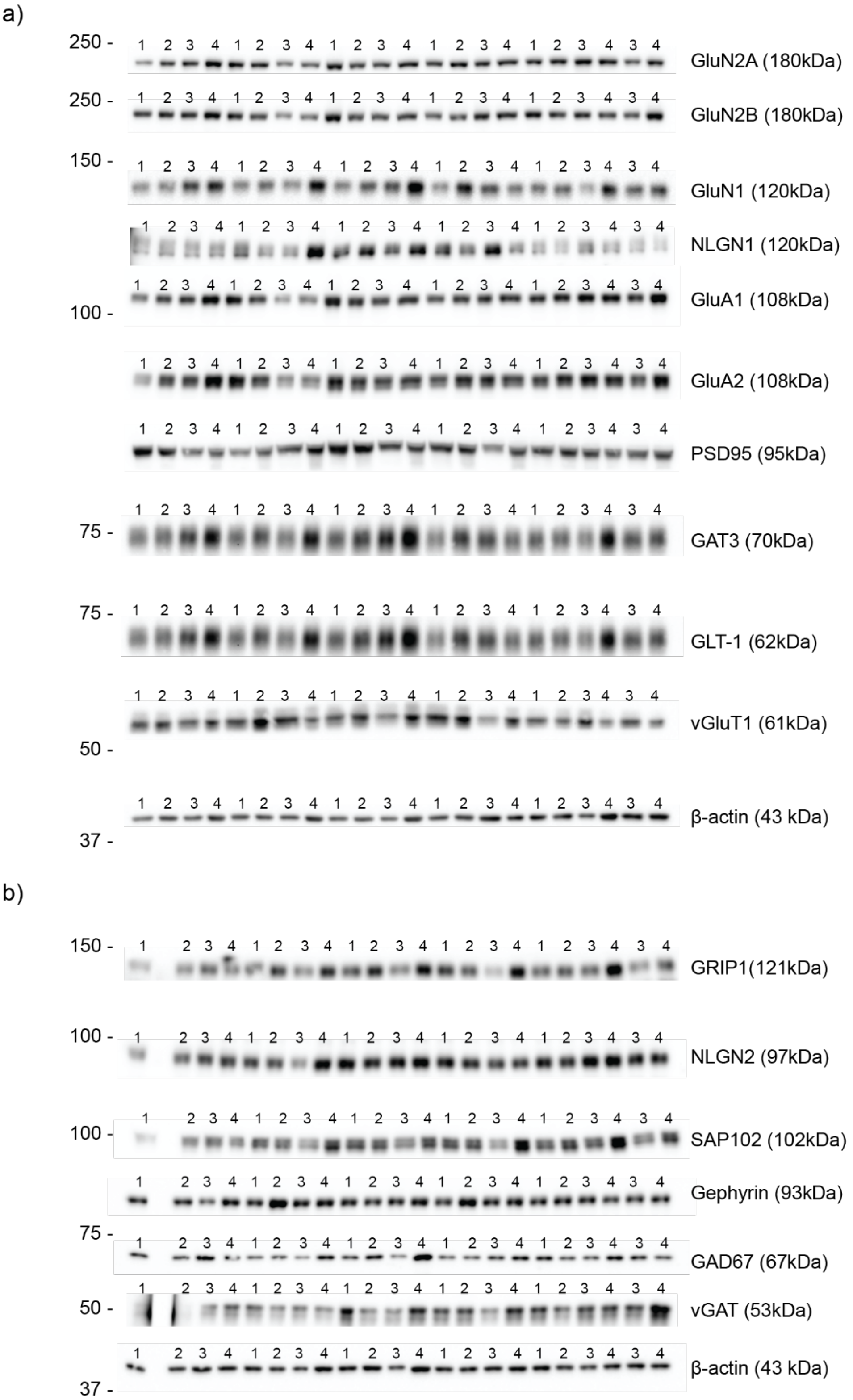

**Supplementary figure 7** Cropped immunoblot related to the protein expression levels of (a) GAT3 (70kDa), GLT-1 (62 kDa), GluA1 (108 kDa), GluA2 (108 kDa), GluN1 (120 kDa), GluN2A (180 kDa), GluN2B (180 kDa), NLGN1 (120 kDa), PSD95 (95 kDa), VGlut1 (61 kDa) and β-actin (43 kDa), and (b) GAD67 (67 kDa), Gephyrin (93 kDa), GRIP1 (121 kDa), NLGN2 (97 kDa), SAP102 (102 kDa), VGAT (53 kDa) and β-actin (43 kDa) measured in the homogenate of the prelimbic cortex of Low SPS naive (1), Low SPS cocaine (2), High SPS naive (3) and High SPS cocaine (4) rats.

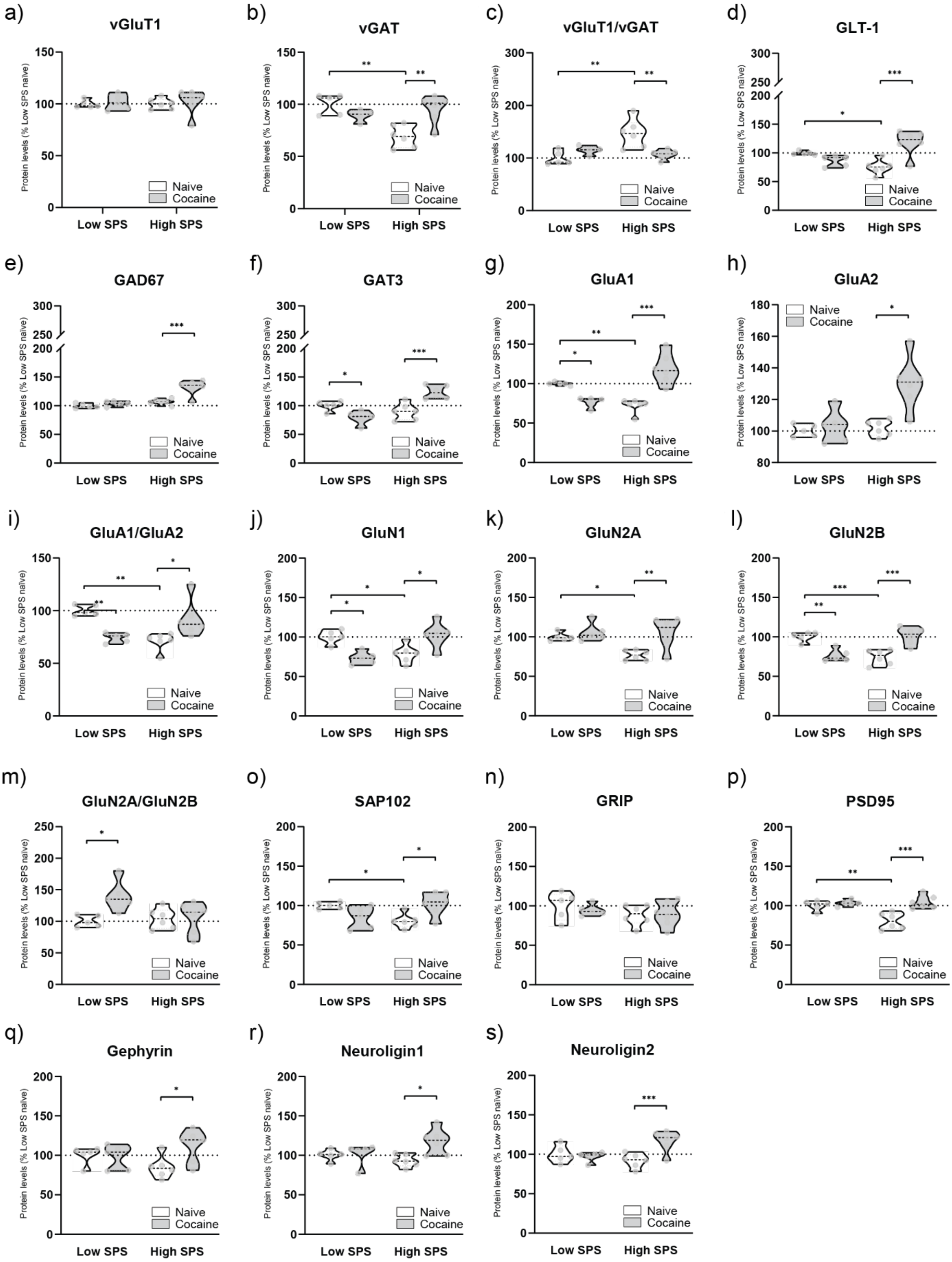

**Supplementary figure 8** The baseline differences between low- and high SPS-like rats (naïve) and cocaine effects in low- and high SPS like rats on glutamatergic- and GABAergic transporters, glutamate-to-GABA converting enzymes, main subunits of AMPA and NMDA glutamate receptors, scaffolding proteins, and structural markers of glutamatergic- and GABAergic synapses in the **infralimbic cortex**. The representative immunoblots can be found in supplementary figure 5. Results are expressed in violin scatter plots as percentage of the mean ± SEM compared to low SPS naïve group (n=5 low SPS naïve, n=5 low SPS cocaine S/A, n=6 high SPS naïve, n=6 high SPS cocaine S/A). Statistics display two-way ANOVA resulted followed by Tukey's post hoc test (*p<0.05, **p<0.01, ***p<0.001) as presented in supplementary table 22, thus without Benjamini-Hochberg correction (supplementary table 23, main body results).

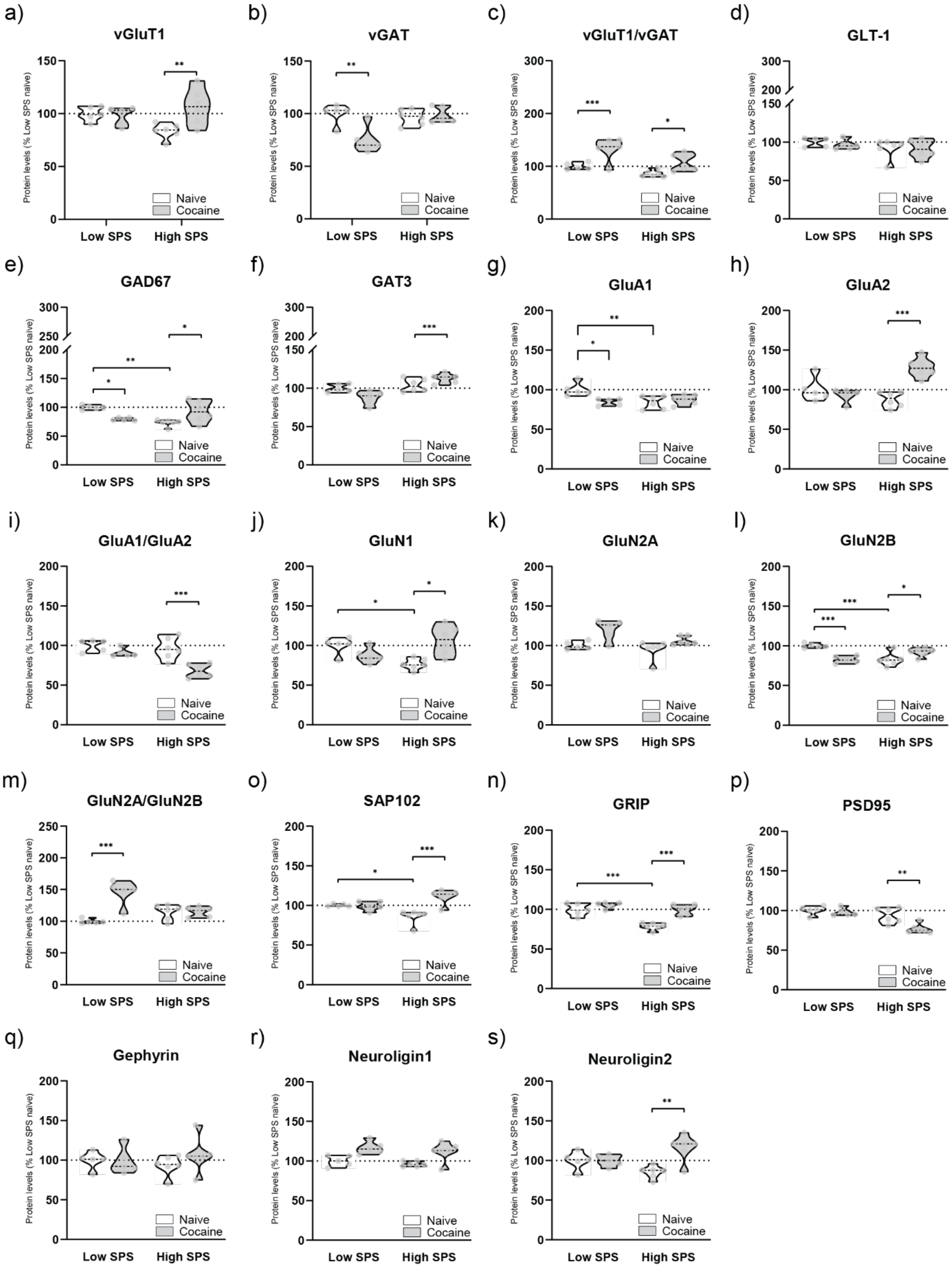

**Supplementary figure 9** The baseline differences between low- and high SPS-like rats (naïve) and cocaine effects in low- and high SPS like rats on glutamatergic- and GABAergic transporters, glutamate-to-GABA converting enzymes, main subunits of AMPA and NMDA glutamate receptors, scaffolding proteins, and structural markers of glutamatergic- and GABAergic synapses in the **prelimbic cortex**. The representative immunoblots can be found in supplementary figure 5. Results are expressed in violin scatter plots as percentage of the mean ± SEM compared to low SPS naïve group (n=5 low SPS naïve, n=5 low SPS cocaine S/A, n=6 high SPS naïve, n=6 high SPS cocaine S/A). Statistics display two-way ANOVA resulted followed by Tukey’s post hoc test (*$p<0.05$, **$p<0.01$, ***$p<0.001$) as presented in supplementary table 22, thus without Benjamini-Hochberg correction (supplementary table 23, main body results).

19. RCoreTeam. R: A language and environment for statistical computing. 2021.

20. How the Dunnett T3, Games and Howell, and Tamhane T2 tests work [GraphPad Statistics Guide]. Available from: https://www.graphpad.com/guides/prism/latest/statistics/stat_multiple-comparisons-without-a.htm.

21. Todorov V. Robust selection of variables in linear discriminant analysis. Statistical Methods and Applications. 2007;15(3):395-407.

22. Todorov V, Filzmoser P. An Object-Oriented Framework for Robust Multivariate Analysis. Journal of Statistical Software. 2009;32(3).

23. West BT, Welch KB, Galecki AT. Linear Mixed Models. A Practical Guide Using Statistical Software2014.

24. Singer JD, Willett JB. Applied Longitudinal Data Analysis: Modeling change and event occurrence.2003.

25. Gueorguieva R, Krystal JH. Move Over ANOVA: progress in analyzing repeated-measures data and its reflection in papers published in the Archives of General Psychiatry. Archives of General Psychiatry. 2004;61(3).

26. Fitzmaurice GM, Laird NM, Ware JH. Applied longitudinal analysis. 2nd ed. Hoboken, N.J.: Wiley; 2011. xxv, 701 p. p.
